# A 3D in vitro assay to study combined immune cell infiltration and cytotoxicity

**DOI:** 10.1101/2024.03.27.586980

**Authors:** Ashleigh J. Crawford, Adrian Johnston, Wenxuan Du, Eban A. Hanna, David Schell, Zeqi Wan, Ting-Hsi Chen, Fan Wu, Kehan Ren, Yeongseo Lim, Praful Nair, Denis Wirtz

**Author notes:** Indicates authors contributed equally (Ashleigh J. Crawford, Adrian Johnston, Wenxuan Du, Eban A. Hanna, David Schell).

## Abstract

Immune cell-mediated killing of cancer cells in a solid tumor is prefaced by a multi-step infiltration cascade of invasion, directed migration, and cytotoxic activities. In particular, immune cells must invade and migrate through a series of different extracellular matrix (ECM) boundaries and domains before reaching and killing their target tumor cells. These infiltration events are a central challenge to the clinical success of CAR T cells against solid tumors. The current standard in vitro cell killing assays measure cell cytotoxicity in an obstacle-free, two-dimensional (2D) microenvironment, which precludes the study of 3D immune cell-ECM interactions. Here, we present a 3D combined infiltration/cytotoxicity assay based on an oil-in-water microtechnology. This assay measures stromal invasion following extravasation, migration through the stromal matrix, and invasion of the solid tumor in addition to cell killing. We compare this 3D cytotoxicity assay to the benchmark 2D assay through tumor assembloid cocultures with immune cells and engineered immune cells. This assay is amenable to an array of imaging techniques, which allows direct observation and quantification of each stage of infiltration in different immune and oncological contexts. We establish the 3D infiltration/cytotoxicity assay as an important tool for the mechanistic study of immune cell interactions with the tumor microenvironment.

**Graphical Abstract:** The 3D combined infiltration/cytotoxicity assay captures three important steps of immune cell infiltration into the solid tumor microenvironment: (1) circulating immune cells extravasate and invade the stromal matrix, (2) immune cells migrate through the stromal matrix to reach the tumor core, and (3) immune cells that successfully navigate the stroma must cross a basement membrane boundary secreted by the cancer cells to contact and kill the cancer cells within a solid tumor.

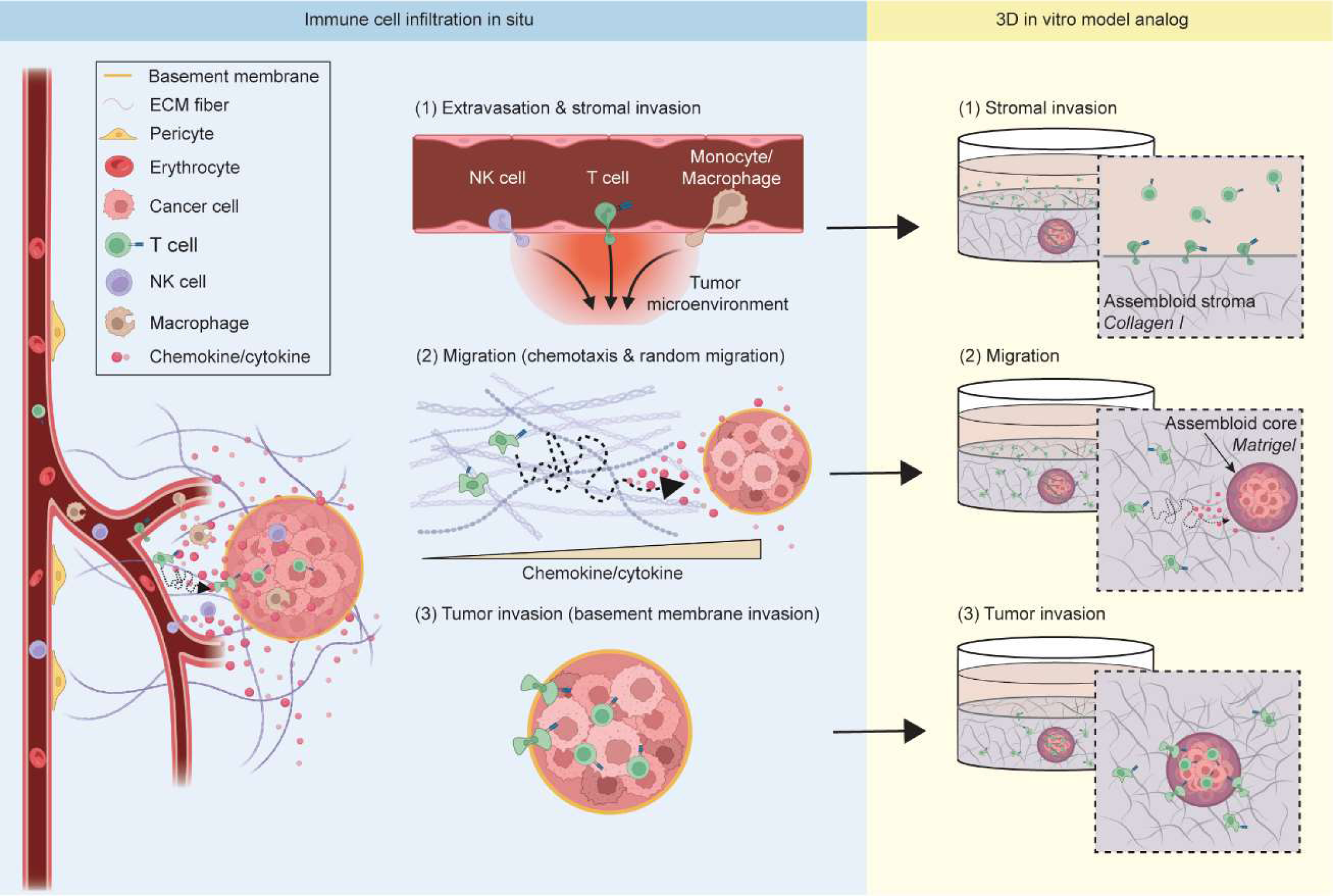

## 1. Introduction

Cytotoxic adaptive (T cells) and innate (macrophages, NK cells) immune cells are recruited to the tumor site for specialized anti-tumor roles in the early stages of tumorigenesis, but their infiltration into the solid tumor microenvironment (TME) is a complex multistep process [1–4]. First, immune cells extravasate from neighboring blood vessels and invade the tumor stroma (Fig. 1a, 1). Next, these cells migrate through the tissue’s stromal matrix toward tumor cells via chemokine/cytokine directed chemotaxis and/or random migration [5] (Fig. 1a, 2). Immune cells then cross the tumor’s basement membrane where they can access cancer cells [6–8] (Fig. 1a, 3). The cytotoxic immune cells that successfully infiltrate the tumor can then kill cancer cells.

**Figure 1.**
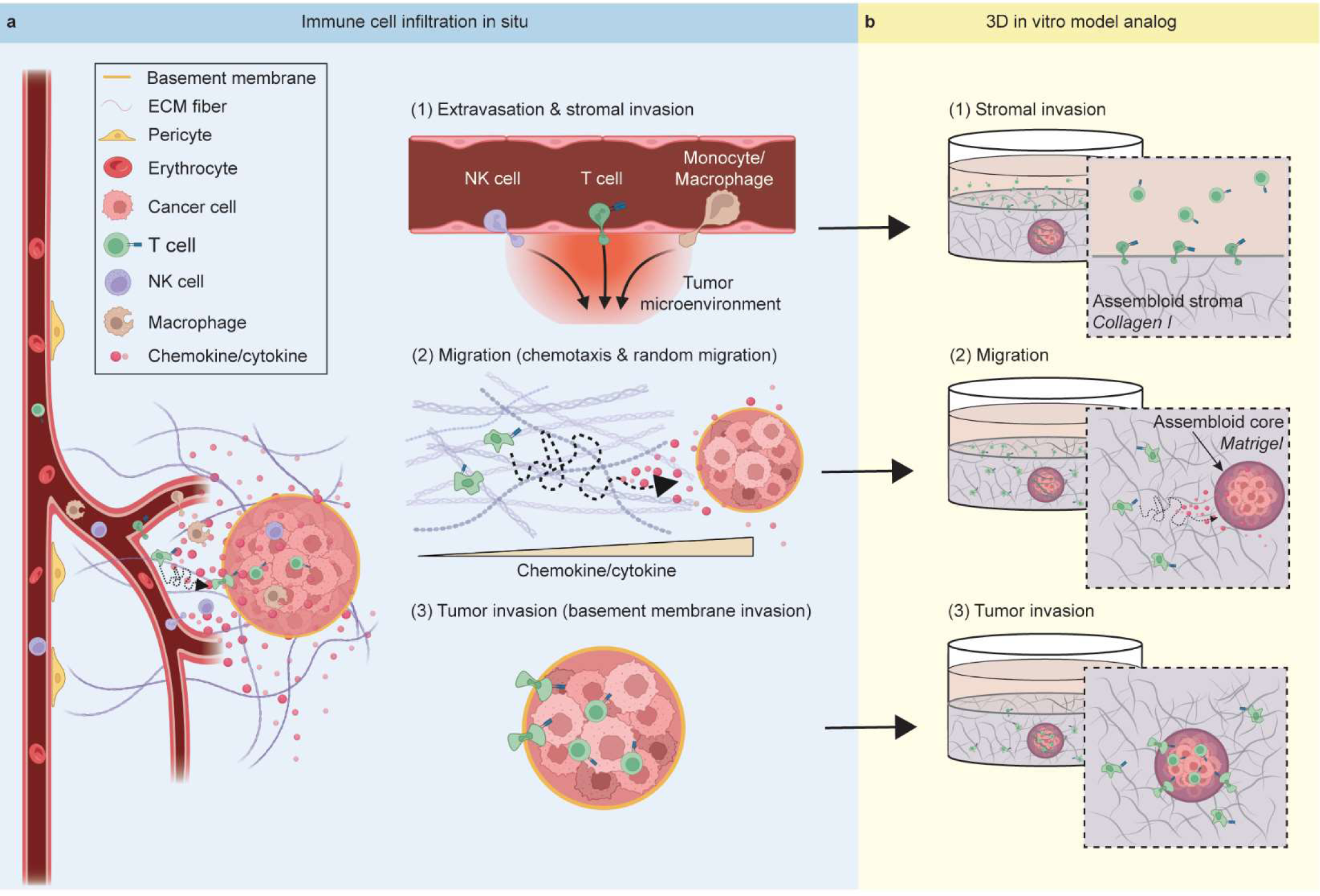
The immune cell infiltration steps in the 3D combined infiltration/cytotoxicity assay are analogous to the infiltration steps observed in situ. (a) In situ events involved in immune cell killing of cancer cells begin immediately following (1) cell extravasation from a blood vessel as the immune cells invade into the tumor stroma. (2) Cell migration through the stromal matrix is driven by chemotaxis along a chemokine/cytokine gradient toward the tumor, or random migration. (3) After reaching the tumor, immune cells need to cross a basement membrane boundary. Once in close proximity or direct contact with cancer cells, immune cells can elicit a cytotoxic effect. (b) In the 3D cytotoxicity assay analog, cancer cells are suspended in a basement membrane forming the assembloid core. This core is embedded in a bulk collagen matrix to simulate the collagen-rich stroma. (1) Immune cells are added in culture medium deposited on top of the assembloid’s bulk matrix. These immune cells must first invade the collagen matrix through the medium-ECM interface. (2) Cells must then migrate to the tumor assembloid core via random migration or chemotaxis following cancer cell secreted chemokines/cytokines. (3) Once reaching the boundary of the assembloid core, immune cells must cross the bulk ECM-assembloid core interface. At this interface, cells leave a collagen rich assembloid stroma to invade the assembloid core, a sphere of basement membrane containing cancer cells. When in close proximity or direct contact with the tumor assembloid core, immune cells can elicit a cytotoxic effector function. To kill cancer cells, immune cells must successfully perform each of these steps in the immune cell infiltration cascade in addition to the cell recognition and killing traditionally tested in 2D cytotoxicity assays.

Current two-dimensional (2D) cytotoxicity assays place immune cells in direct contact with their target cancer cells, bypassing these important infiltration steps. These assays focus exclusively on proximity-mediated cell recognition and killing by co-seeding immune cells with cancer cells or overlaying immune cells onto a monolayer of cancer cells on a culture plate [9–11]. Yet, the ability of immune cells to travel to the tumor site is an essential cellular attribute when evaluating their overall cytotoxic effect in solid tumors [4,5,12–14]. The physical barriers dividing the collagen-rich stromal region from the tumor where cancer cells deposit the components of a basement membrane (i.e., laminins and collagens) cannot be replicated in 2D systems [6–8,15]. Moreover, cell phenotypes are altered in 2D compared to 3D environments [16,17]. Gravity pulls cells to the bottom of the culture plate, which forces artificially enhanced interactions between immune and cancer cells in 2D. Highly dynamic focal adhesions formed by cells interacting with the extracellular matrix (ECM) and non-apical cell polarization play key roles in cell migration; they are some features of the 3D TME that cannot be recapitulated in 2D systems [16–18]. Cell secretory profiles are also altered [19], leading to artificial or absent chemokine/cytokine gradients in 2D [20]. Stromal cells, including cancer-associated fibroblasts (CAFs) and tumor-associated macrophages (TAMs) [21], secrete these soluble factors in the 3D collagen-rich stromal region adjacent to but spatially distinct from cancer cells [15]. In sum, accurate in vitro modeling of this geometry and the physiological cell interactions with the TME requires advanced 3D systems.

While some 3D spheroid coculture models exist to study immune-cancer interactions [22–24], these systems premix cancer and immune cells in one ECM (commonly Matrigel, a solubilized basement membrane) or medium, presenting an artificial solid tumor microenvironment with non-physiological cell-ECM interactions that still bypass the infiltration steps required of immune cells. To help study these important immune cell infiltration events and interactions with the ECM, we developed a 3D combined infiltration/cytotoxicity assay that captures the multistep process of infiltration into a solid tumor (Fig. 1b). This in vitro system uses an oil-in-water microtechnology [8] to define spatially distinct tumor core and tissue stroma regions within an assembloid, a 3D in vitro system that cocultures multiple cell types in adjacent compartments resembling the in situ tissue architecture [15,25,26]. In our multi-compartment assembloid of the tumor microenvironment, cancer cells and immune cells are cultured in adjacent compartments where cell and ECM compositions are tuned to mimic the tissue composition of a solid tumor [15]. We vary the location where effector cells are seeded with respect to cancer cells to simulate and study each key immune cell infiltration event and measure the infiltration and cytotoxicity of human T cells, macrophages, and natural killer (NK) cells. All infiltration events are captured in the final assembloid geometry: immune cells initially placed in cell culture medium invade the collagen matrix, mimicking the stromal invasion that follows immune cell extravasation (Fig. 1b, 1). These immune cells then migrate through the collagen matrix toward the tumor assembloid core following chemotactic gradients or random migration (Fig. 1b, 2). Once immune cells arrive at the assembloid core, these cells must cross a basement membrane boundary to be in direct contact with or close proximity to cancer cells (Fig. 1b, 3). Thus, our 3D in vitro analog simulates the motility and invasion steps preceding immune cell killing, after which the infiltrated immune cells can elicit an effector function which we measure via their cytotoxicity against the cancer cells in the assembloid core. This 3D infiltration/cytotoxicity assay can be used to study immune cell interactions within the stromal compartment and its cellular components as well, such as how the 3D TME can both support and restrict immune cell functions [27].

Such insights would not only advance our mechanistic understanding of immune cell motility in the TME, but also address a significant knowledge gap in engineered cell therapies [28–30]. These cell therapies, including chimeric antigen receptor T (CAR T) cells, have been successful against hematological malignancies where engineered cells are intravenously (IV) injected into direct contact with cancer cells in the blood [4,28,30–33]. However, CAR T cell therapies have been unsuccessful thus far as a treatment for solid tumors where navigation of the surrounding TME is required before contact with cancer cells [28–30,34]. Our recent work addresses this central challenge with current CAR T cells: their poor infiltration in solid tumors [4]. Standard CAR T cells typically remain confined to the edges of the solid tumor, precluding cancer-cell killing, which has led to their limited success in recent clinical trials [28–30,34]. Our approach consists of engineering self-propelled tumor-infiltrating CAR T cells by expressing synthetic “velocity receptors” [4]. The design and development of other CAR T cells with enhanced tumor-infiltration properties require a new 3D assay that acknowledges that in order to kill target cancer cells, CAR T cells need to efficiently extravasate, migrate through the collagen-rich stromal matrix, and cross the basement membrane confining tumor cells. Here, we demonstrate our 3D assay’s versatility when studying these events by evaluating the infiltration and cytotoxicity of different immune cells (macrophages, T cells, NK cells, and CAR T cells) targeting different types of cancer cells, different spatial distributions of immune cells relative to cancer cells to capture key immune infiltration events, extended experimental time durations, and coculture capabilities with different types of stromal cells. Our 3D assay can be used to study and enhance the invasion and migration capabilities of engineered cells [35] and improve the prediction of in vivo outcomes [4], thus such 3D tools are critically important to assessing immune cell infiltration of the TME.

## 2. Materials and Methods

### 2.1 PBMC isolation

Primary human T cells, monocytes, and NK cells were isolated and purified from peripheral blood mononuclear cells (PBMCs) of buffy coats obtained from healthy volunteers (Blood donation center, Anne Arundel Medical Center, Annapolis, MD, USA) using Ficoll-Hypaque (Ficoll Paque Plus, Sigma, GE17-1440-02) according to the manufacturer’s instructions. Briefly, blood was diluted in RPMI 1640 (Gibco 11875-093), and the diluted blood was overlaid onto Ficoll-Hypaque. Plasma, PBMCs, Ficoll-Hypaque, and red blood cell layers were separated after centrifugation (30 min, 400 x g), and the PBMC layer was harvested and centrifuged (10 min, 400 x g). PBMCs were resuspended in freezing medium (90% FBS [Corning 35-010-CV] + 10% Dimethyl sulfoxide [DMSO, ATCC 4-X]) at a concentration of 5 x 10^7^ cells/mL. Cryovials containing PBMCs were initially frozen at −80°C and transferred into liquid nitrogen for long-term storage.

### 2.2 T cell purification and CAR T generation

Primary human CD4 and CD8 T cells were isolated from PBMCs by negative selection using STEMCELL EasySep isolation kits (Stemcell Technologies, 17952 and 17953), and mixed 1:1. T cells were then activated overnight by CD3/CD28 Dynabeads (Gibco, 11131D) with 100 IU/ml IL2 (Peprotech, 200-02) in X-Vivo 15 media (Lonza, BE02-053Q) supplemented with 5% human AB serum (Millipore Sigma, H4522), 2 mM L-glutamine (Gibco, 25030081), and 100 U/ml penicillin and 100 mg/ml streptomycin (Gibco, 15140122). After overnight activation, T cells were transduced using lentivirus at an MOI between 5 and 10 on Retronectin-coated (Takara, T100B) plates overnight to express an M5 or SS1 CAR. The M5 or SS1 CAR [36,37] was inserted into the backbone of the pSLCAR-CD19-BBz plasmid (Addgene 135992) by standard molecular cloning techniques using Gibson HiFi DNA assembly (NEB, E2621L) to lead to the lentiviral integration of an M5 or SS1 CAR – P2A – eGFP. M5 or SS1 CAR lentiviral particles were generated by transfecting 293T cells using GeneJuice (Millipore Sigma, 70967). Lenti-X Concentrator (Takara, 631232) was used to concentrate lentivirus, after which the lentivirus concentration was measured by p24 ELISA (Takara, 632200). Following overnight transduction, CAR positivity was measured by GFP expression through flow cytometry. CAR positivity was routinely greater than 75% positive. CAR T cells and non-transduced T cells were then expanded for 9 – 12 days in media and 100 IU/ml IL2 before use in experiments or storage in liquid nitrogen.

### 2.3 Macrophage purification

Macrophage attachment medium was prepared using DMEM (Corning 10-013-CV) supplemented with 10% heat-inactivated (57 °C for 30 min) FBS (Corning 35-010-CV), 1% penicillin-streptomycin (Sigma P0781) and 50 ng/mL recombinant human M-CSF (R&D Systems 216-MC). M1 macrophage attachment medium was prepared using DMEM supplemented with 10% heat-inactivated FBS, 1% penicillin-streptomycin and 50 ng/mL recombinant human GM-CSF (R&D Systems 215-MC). M1 macrophage differentiation medium was prepared using M1 macrophage attachment medium supplemented with 100 ng/mL of LPS (Sigma-Aldrich L2630) and 50 ng/mL of IFN-γ (R&D Systems 285-IF).

Primary human monocytes were isolated from PBMCs from three donors using a negative selection classical monocyte isolation kit (Miltenyi Biotec 130-117-337) according to the manufacturer’s protocol. Isolated monocytes were counted using 3% acetic acid with methylene blue solution (STEMCELL 07060) to identify and lyse residual red and while red blood cells. Monocytes were seeded in a 6-well tissue culture plate (Corning 353046) at 2.5 x 10^5^ cells/mL in macrophage attachment medium. After three days, monocytes that remained in the suspension were gently aspirated away, the attached macrophages were washed three times with DPBS (Corning 21-031-CV), and fresh macrophage attachment medium was added for M0 naïve macrophages. On day 5, fresh macrophage attachment medium was added without wash for M0 naïve macrophages. For M1 macrophage priming, M1 macrophage attachment medium was added on day 3 instead. On day 5, M1 macrophage differentiation medium was added to M1 macrophages after 3 times wash with DPBS. On day 7, both M0 naïve macrophages and M1 macrophages were treated with pre-warmed TrypLE^TM^ Express Enzyme (Gibco 12604013) for 10-15 min and were gently detached by pipetting. TAMs were differentiated from M0 naïve macrophages by stimulation with PANC-1 conditioned medium for 48 h and confirmed by flow cytometry (Supplementary Fig. 5). TAMs were stained with CellTracker^TM^ Red CMTPX Dye (Invitrogen^TM^ C34552) diluted 1:1000 in serum free DMEM (Corning 10-013-CV) for 30 min before detaching with pre-warmed TrypLE^TM^ Express Enzyme (Gibco 12604013) for 10-15 min and gently pipetting. Harvested macrophages were kept on ice before seeding for the 2D/3D killing assay. Cells were maintained at 37 °C, 5% CO_2_ in a humidified incubator.

### 2.4 Flow cytometry

Different phenotypes of macrophages were detached via TrypLE^TM^ Express Enzyme (Gibco 12604013) for 10-15 minutes in an incubator (37 °C, 5% CO_2_) and washed 3 times in DPBS before resuspending at a concentration of 1 x 10^6^ cells/mL in the FACS (fluorescence-activated cell sorting) wash buffer (1x DPBS [Corning 21-031-CV] + 2% FBS [Corning 35-010-CV] + 0.1% NaN_3_ [Sigma SX0299]). Cell suspensions were blocked with Human TruStain FcX (Biolegend 422301) for 15 min under room temperature. The antibody staining solution was then added directly (5µL per million cells) and cell suspensions were incubated at 4 °C for 30 min, protected from light. Conjugated antibodies used for cell labeling were as follows: APC anti-human CD80 (Biolegend 375403), PE anti-human CD163 (Biolegend 333605), PE/Cyanine7 anti-human CD206 (Biolegend 321123). Unstained control samples were prepared via staining with APC Rat igG2a, κ Isotype Ctrl (Biolegend 400511), PE Mouse IgG1, κ Isotype Ctrl (Biolegend 400111), PE/Cyanine7 Mouse IgG1, κ Isotype Ctrl (Biolegend 400125). After staining, cell samples were washed in 3mL of FACS wash buffer and resuspended at a concentration of 1 x 10^6^ cells/mL in FACS wash buffer. Samples were kept on ice until they were analyzed on the BD FACSCanto^TM^ flow cytometry system. Data analysis was performed using FlowJo software.

### 2.5 NK cell purification

Primary human NK cells were isolated from PBMCs from three donors by negative selection using the EasySep^TM^ Human NK Cell Isolation Kit (STEMCELL Technologies 17955), according to the manufacturer’s protocol. Immediately following isolation, NK cells were expanded using the ImmunoCult^TM^ NK Cell Expansion Kit (STEMCELL Technologies 100-0711), per the manufacturer’s protocol. After expansion for 14 days, NK cells were harvested for seeding in the 2D/3D killing assays. Prior to coculture in the PDAC assembloid system, NK cells were stained with CellTracker^TM^ Green CMFDA (5-chloromethylfluorescein diacetate) (Invitrogen^TM^ C7025) diluted to 10 µM in serum free ImmunoCult^TM^ NK Cell Base Medium (STEMCELL Technologies 100-0712) for 30 min. Cells were maintained at 37 °C, 5% CO_2_ in a humidified incubator.

#### Cell culture

OVCAR3 ovarian cancer cells (ATCC HTB-161), MDA-MB-231 breast cancer cells (ATCC HTB-26), JAR gestational trophoblastic neoplasia cells (ATCC HTB-144), and PANC-1 pancreatic ductal adenocarcinoma cells (ATCC CRL-1469) were cultured according to ATCC recommendations. OVCAR3 cells were cultured in RPMI 1640 (Gibco 11875-093) supplemented with 10% FBS (Corning 35-010-CV), 1% sodium pyruvate (Sigma S8636), 1% HEPES (Gibco 15630-080), 1% glucose (Gibco A24940-01), and 1% penicillin-streptomycin (Sigma P0781). JAR cells were cultured in RPMI 1640 (Gibco 11875-093) supplemented with 10% FBS (Corning 35-010-CV) and 1% penicillin-streptomycin (Sigma P0781). MDA-MB-231 cells, PANC-1 cells and pancreatic ductal adenocarcinoma cancer-associated fibroblasts (CAF-SC2) were cultured in DMEM (Corning 10-013-CV), supplemented with 10% FBS (Corning 35-010-CV), and 1% penicillin-streptomycin (Sigma P0781). Cell passages did not exceed 20. Cells were maintained at 37 °C, 5% CO_2_ in a humidified incubator.

### 2.6 Lentiviral transduction/luciferase tag

The 293T (ATCC CRL-3216) immortalized cell line was cultured with DMEM (Corning 10-013-CV), supplemented with 10% FBS (Corning 35-010-CV), and 1% penicillin-streptomycin (Sigma P0781). Cell passages did not exceed 20. Cells were maintained at 37 °C, 5% CO_2_ in a humidified incubator.

To generate the OVCAR3 and JAR luciferase-expressing mCherry-tagged cell lines, the luciferase (Addgene 46793) and mCherry (Addgene 84020) sequences were inserted into the pSLCAR-CD19-BBz (Addgene 135992) plasmid via molecular cloning. The plasmid region from the promoter to the end of the GFP sequence was removed from the original plasmid.

293T cells were transfected with the modified plasmid using GeneJuice (Novagen 70967) mixed with a combination of the assembled plasmid containing the luciferase tag, PsPAX2 (Addgene 12260), and VSV-G (Addgene 12259). ViralBoost (Alstembio VB100) and Opti-MEM (Gibco 31985-070) supplemented with 5% FBS (Corning 35-010-CV) were used to increase viral production. The cells were maintained at 37 °C, 5% CO_2_ in a humidified incubator for 3 days. The virus was then centrifuged at 500 x g for 10 min and filtered through 0.45 um filter to remove 293T cells. Lenti-X Concentrator (Takara Bio PT4421-2) was added at a 1:3 volume ratio with clarified supernatant (1 part Lenti-X Concentration: 3 parts lentivirus-containing supernatant). The mixture was incubated at 4 °C overnight and then centrifuged at 1,500 x g for 45 min at 4 °C the following day. The supernatant was removed, and the pellets were resuspended in 2% of the original volume in cancer cell culture medium.

Cancer cells were transduced with RetroNectin reagent (Takara Bio T100B) according to the manufacturer’s protocol. In brief, the RetroNectin reagent was diluted with DPBS (Corning 21-031-CV) to 40 µg/mL and added to a 6-well plate. The plate was stored at 4 °C overnight. The next day, the plate was blocked with 2 mL of 2% bovine serum albumin (Sigma Aldrich 9048-46-8) in DPBS and washed with 2 mL of HBSS (Sigma Aldrich H6648). The luciferase-mCherry virus (1.2 mL) was added immediately to the plate, and after 6 h of incubation at 37 °C in a 5% CO_2_ humidified incubator, OVCAR3 cells were added. The cells and virus were co-incubated for 3 days. Transduced mCherry-positive cells were sorted by fluorescence-activated cell sorting (FACS, >95%) with a SH800S Cell Sorter (Sony Biotechnology).

### 2.7 Specific lysis calculation

Specific lysis was calculated according to Equation (1) [38,39] below. Cancer cells were luciferase expressing and immune cells did not express luciferase so that cancer cell relative proliferation can be measured independently from immune cells in coculture. The cancer cell luciferase activity was measured for control, dead (1% Triton-X-100 [Sigma T9284] for 24 h), and experimental condition wells. The control_avg_ value is the baseline proliferation of cancer cells in the assembloid, the dead_avg_ value is the background value for dead cells with no luciferase activity, and the specific lysis (%) value calculated for each experimental condition represents where the experimental condition falls within the range of luciferase activity for live control wells (0% specific lysis) to dead (100% specific lysis). In other words, specific lysis calculates the percentage of cancer cells that were lysed by immune cells.

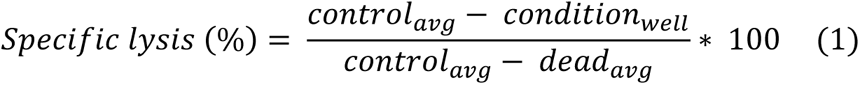

### 2.8 2D cytotoxicity assay

Luciferase-tagged cancer cells were seeded at 5 x 10^3^ cells/well in a 96-well white polystyrene microplate (Corning 3903) overnight. Immune cells at the indicated effector to target (E:T) ratios were added and co-cultured with OVCAR3 cells at 37 °C, 5% CO_2_ in a humidified incubator for 48 h. Triton-X-100 (Sigma T9284) was added (final concentration 1%) to the positive control wells (dead), whereas immune cell media was added to the negative control wells (live control). Then, 100 µL of the supernatant was removed from each well, and 100uL of Bright-Glo luciferase assay reagent (Promega PRE2620) was added according to the manufacturer’s instructions. The plate was covered with foil to prevent light exposure and shaken on an ORBI-SHAKER (Benchmark) at 400rpm for 5 min. The plate was left at room temperature for another 7 min before luminescence (integration time 500ms and second reading 1000ms) was read on a SpectraMax plate reader (Molecular Devices). Specific lysis was calculated using Equation (1).

### 2.9 3D cytotoxicity assay

The 3D cytotoxicity assay uses an oil-in-water microtechnology to generate multi-compartment tumor assembloids as we have previously described [8,40,41]. Briefly, cells (luciferase-tagged cancer cells with immune cells in the core coculture condition or without in all other conditions) are mixed in Matrigel at 2 x 10^4^ cells/µL. High concentration, LDEV-free Matrigel (Corning 354248) was diluted 1:1 with serum free DMEM (Corning 10-013-CV) before mixing with cancer cells. Droplets (2 µL) of the Matrigel/cell mixture were pipetted into mineral oil columns and allowed to gel at 37 °C, 5% CO_2_ incubator for 1 h. These assembloid cores were collected in warmed serum free DMEM and the mineral oil washed off. One core was embedded in 100 µL of 2 mg/mL collagen I (Corning 354249) per well of a 96 well plate (Corning 3903 or Cellvis P96-1.5H-N). Immune cells were mixed in the collagen mixture in the matrix coculture condition High concentration rat tail collagen I was diluted to 2 mg/mL as we have previously described [8,40–43] by mixing culture medium, RECON buffer (1.1 g sodium bicarbonate (Sigma S5761) and 2.4 g HEPES (Acros organics CAS 7365-45-9) in 50 ml milli-Q water), collagen I, and 1M NaOH (Sigma 137031). After a 1.5 h incubation at 37 °C, 5% CO_2_, 100 µL of immune cell culture medium was added on top of the gel. In the medium coculture condition, immune cells were included in the added medium. E:T ratios of 1:1, 1:5, 1:10 were used for the immune and cancer cells. Cancer assembloids were cocultured with immune cells for 2, 4, or 6 days before endpoint luciferase assays were performed.

A collagenase solution was prepared at 4 mg/mL (Gibco 17104019 or 17018029) in DPBS (Corning 21-031-CV). The medium from each well was removed and each well was washed twice with 100µL of pre-warmed DPBS. Then, 50 µL of collagenase solution was added to each assembloid and incubated for 1h (37°C, 5% CO_2_ in a humidified incubator). After the incubation, we added 150 µL of a BrightGlo-TritonX solution (90% BrightGlo + 1% TritonX + 9% DPBS) to each well. The plate was then wrapped in aluminum foil and put on the shaker plate for 5 min at 400 rpm. The plate was then left at room temperature for 7 min before reading luminescence according to the BrightGlo manufacturer’s instructions (Promega PRE2620, integration time 500ms and second reading 1000ms) on a SpectraMax plate reader (Molecular Devices). Specific lysis was calculated using Equation (1).

### 2.10 Assembloid imaging

Assembloids were imaged using phase-contrast, epifluorescence, differential interference contrast (DIC), and fluorescence confocal microscopy of live assembloids. Phase-contrast and epifluorescence images were obtained on a Nikon Ti2 inverted microscope with a DS-Qi2 camera, a Tokai Hit stage top incubator with temperature and CO_2_ control and a 10X/0.30 Plan Fluor Ph1 DL objective (Nikon). DIC and confocal microscopy of live assembloids were obtained on a Nikon A1R confocal/Nikon Eclipse Ti inverted microscope or a Leica Stellaris-5 confocal microscope. The Nikon confocal microscope was equipped with an OKO Labs stage-top incubator with temperature and CO_2_ control, a 10X/0.30 Plan Fluor objective (Nikon), a N1 DIC condenser (Nikon), and a 10X DIC slider (Nikon). The Leica confocal microscope was equipped with a Tokai Hit stage top incubator with temperature and CO_2_ control and a 10X objective (Leica).

### 2.11 Immunofluorescence

Assembloids were fixed in formalin (10%, VWR 16004-121) for 24 hours. Assembloids were then paraffin embedded and the center plane sectioned (4 µm thickness) at Johns Hopkins Oncology Tissue Services (SKCC). Immunofluorescence staining was performed using the Opal fluorescence immunohistochemistry kit according to the manufacturer’s protocol. In short, after deparaffinization and rehydration, sections were blocked with peroxidase and alkaline phosphatase blocking reagent (Dako S200389-2) for 30 minutes. Then, sections were incubated with anti-CD11b primary antibody (Abcam AB133357) overnight. Then, sections were incubated with Opal Anti-Ms + Rb HRP (Akoya Biosciences ARH1001EA) for 20 min, followed by incubation with TSA plus Cyanine 5 (Akoya Biosciences NEL745001KT) for 15 min. Finally, sections are stained with Hoechst 33342 (Thermo Fisher Scientific 62249) for 15 min and mounted in ProLong Diamond Antifade Mountant (Thermo Fisher Scientific P36961). Sections were imaged using a Nikon Swept Field Confocal Microscope at 10× magnification. Thereafter, whole slide images were stitched and cropped with Matlab, and visualized using ImageJ.

### 2.12 Cartoons

Cartoons were generated with Adobe Illustrator and BioRender.com.

### 2.13 Statistical analysis

Statistical analyses are indicated in figure legends with P values included.

## 3. Results

### 3.1 The 3D cytotoxicity assay

Our 3D infiltration/cytotoxicity assay captures the key steps involved in immune cell infiltration of solid tumors (Fig. 1, a and b). This assay utilizes a multi-compartment assembloid technique to encase cancer cells in a basement membrane and embed this tumor assembloid core in a bulk collagen I matrix [8,40–42]. Later, we describe different assembloid geometries that were designed to isolate individual infiltration steps.

Widely used to study immune cell cytotoxicity, the standard in vitro cell killing assay grows cancer cells in 2D on plastic or glass (Fig. 2a) [9–11]. Immune cells are co-seeded with cancer cells or overlayed on top of a cancer cell monolayer (Fig. 2b). This 2D assay tests the ability of immune cells to target cancer cells and their immediate cytotoxicity. For OVCAR3 human ovarian cancer cells, which are commonly used in the assessment of mesothelin-specific CAR T cells [36,37,44,45], the cytotoxicity of M5 CAR T cells was > 90% in the standard 2D cell-cell recognition platform at all tested E:T (Effector:Target) ratios (Fig. 2c). Cytotoxicity was measured using a specific lysis calculation from a standard luciferase assay that measures the relative proliferation of luciferase-expressing cancer cells independent of immune cell proliferation by comparing the experimental condition to luminescence values of live and dead cancer cell monocultures (more information in Methods) [38,39].

**Figure 2.**
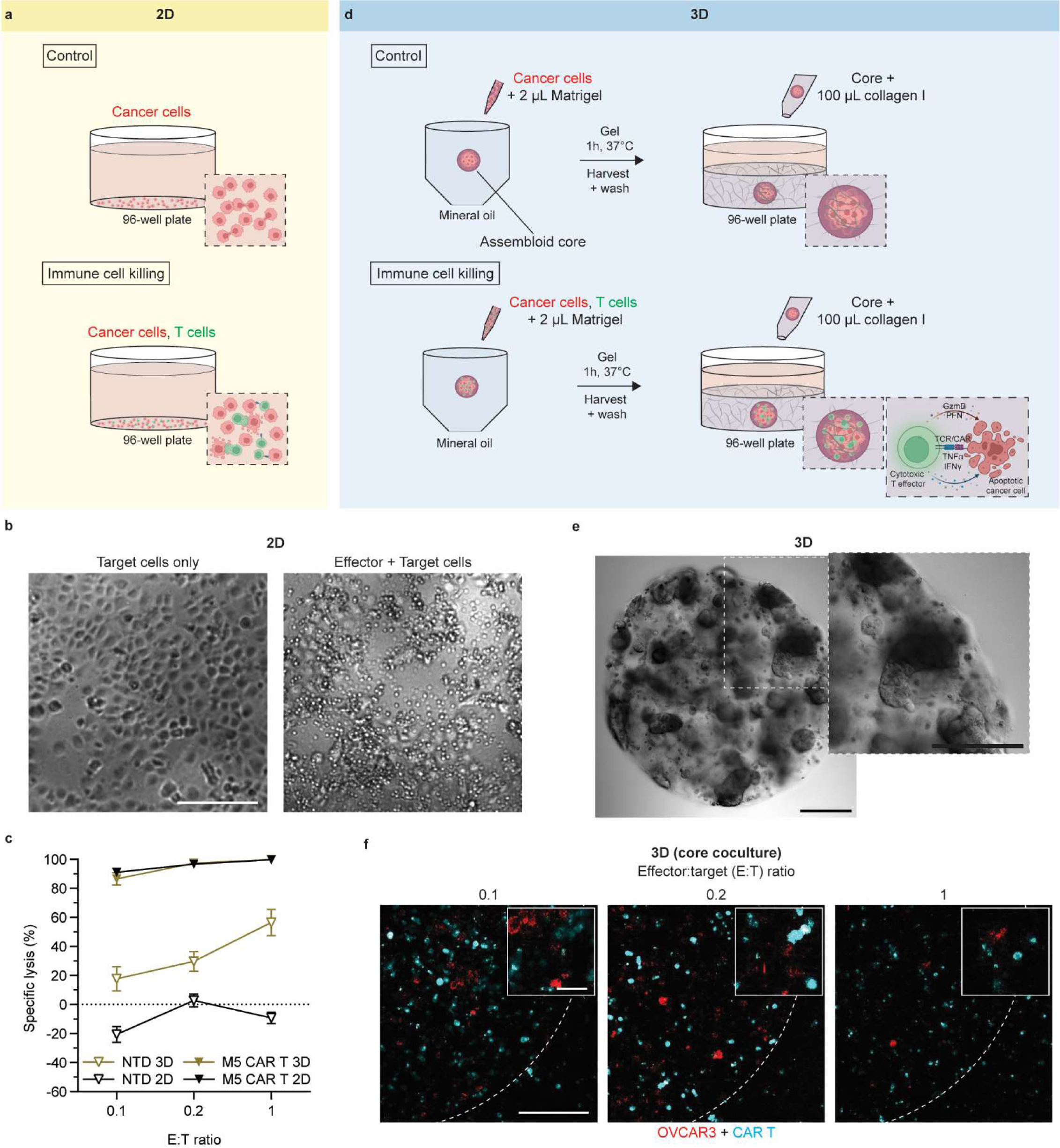
Standard 2D vs. 3D cytotoxicity assay. (a) Cartoon depicting the standard 2D cytotoxicity assay. Cancer cells (red) are in monoculture (control) or coculture (immune cell killing) with immune cells (green) on plastic or glass in a 96-well cell culture plate. (b) Phase-contrast images of the 2D cytotoxicity assay on day 2. Target cancer cells adhered to the bottom of the plate. Effector cells that were premixed with target cells were much smaller in diameter and did not always adhere to the plate. 10X magnification. Scale bar, 250 µm. (c) The cytotoxic effect of M5 CAR T cells and non-transduced (NTD) T cells against OVCAR3 ovarian cancer cells in 2D and 3D on day 2. Specific lysis calculation is described in Methods. E:T, Effector:Target ratio. N (biological replicates) = 3, n (technical replicates) = 3. Data are mean ± SEM. (d) Cartoon depicting the new 3D cytotoxicity assay including the 3D assembloid generation technique. Assembloids are plated in a 96-well plate. Cancer cells or a mixture of cancer cells and immune cells are suspended in small spherical volumes of a basement membrane (Matrigel) generated using oil columns. Assembloid cores are then embedded into a bulk collagen I matrix. The cytotoxic interactions between cancer cells and immune cells are shown. (e) Differential Interference Contrast (DIC) image of an assembloid used in the 3D assay. 10X magnification. Scale bars, 250 µm. (f) Assembloids used in the 3D assay with T cells and cancer cells cocultured in the assembloid core. OVCAR3 cells were labeled with mCherry (red) and M5 CAR T cells were labeled with eGFP (cyan). Dotted lines indicate the boundary of the assembloid cores. Assembloids were imaged on day 2, and images are maximum intensity projections of stacks of confocal fluorescence microscopy images. 10X magnification. Scale bar, 150 µm. Inset, 30 µm.

To simulate in situ cytotoxic activity in a 3D system after immune cells infiltrate the tumor, we used an oil-in-water microtechnology to generate tumor assembloids with cancer cells isolated in a central compartment [8]. Immune cells were cocultured with cancer cells in a confined 2 µL droplet of basement membrane in this first assembloid geometry (Fig. 2d). This assembloid core was embedded in a bulk collagen I matrix (2 mg/mL), designed to simulate the collagen dense stroma surrounding a solid tumor [8,15,41–43] (Fig. 2e); although, other ECM components can be added in this compartment if necessary. We embedded the assembloid core in a 100 µL bulk ECM to ensure the boundary effects of the system on cell migration were negligible [41]. We note that cancer cells primarily migrate in collagen while they primarily proliferate in Matrigel (basement membrane) [8,41,46–50], thus this compartmentalization of ECMs is required for studies of physiological cell-ECM interactions. Importantly, this first version of the assay only measures the cytotoxic effect of immune cells in 3D compared to 2D. This geometry does not evaluate the infiltration requirements that precede these interactions, but rather measures the ability of immune cells to recognize and kill cancer cells after they have invaded the tumor. The 3D cytotoxic effect of M5 CAR T cells was similar to that observed in the 2D assay (Fig. 2c). M5 CAR T cells (targeting) were also significantly more cytotoxic than non-transduced (NTD) T cells (non-targeting) in both 2D and 3D, as expected [9,22]. Interestingly, in this (3D) core coculture system, some but not all M5 CAR T cells seeded in the assembloid core migrated from the core into the surrounding collagen matrix (Fig. 2f, Supplementary Fig. 1a left panel). This observation demonstrates an additional application of this assay in studies investigating immune cell persistence at the tumor site.

### 3.2 The multi-compartment assembloid simulates the critical steps of immune cell infiltration

The 3D assembloid system is a dual-compartment model defined by two ECMs - basement membrane (tumor) and collagen I (stroma) - which are the two predominant ECMs in the tumor microenvironment [15,51]. The immune, stromal, and cancer cell compositions of each compartment can be customized. We measured the infiltration and cytotoxicity of NTD and M5 CAR T cells in three different assembloid geometries (Fig. 3a). Each geometry was designed to progressively add immune-TME navigation steps until all key steps of immune cell infiltration were simulated (Fig. 1, Fig. 3b). The OVCAR3 cancer cell count of each geometry was compared to the OVCAR3 cell counts of control assembloid monocultures (Fig. 3a top panel – control, Supplementary Fig. 2a) after two days of assembloid culture using a specific lysis calculation (more information in Methods).

**Figure 3.**
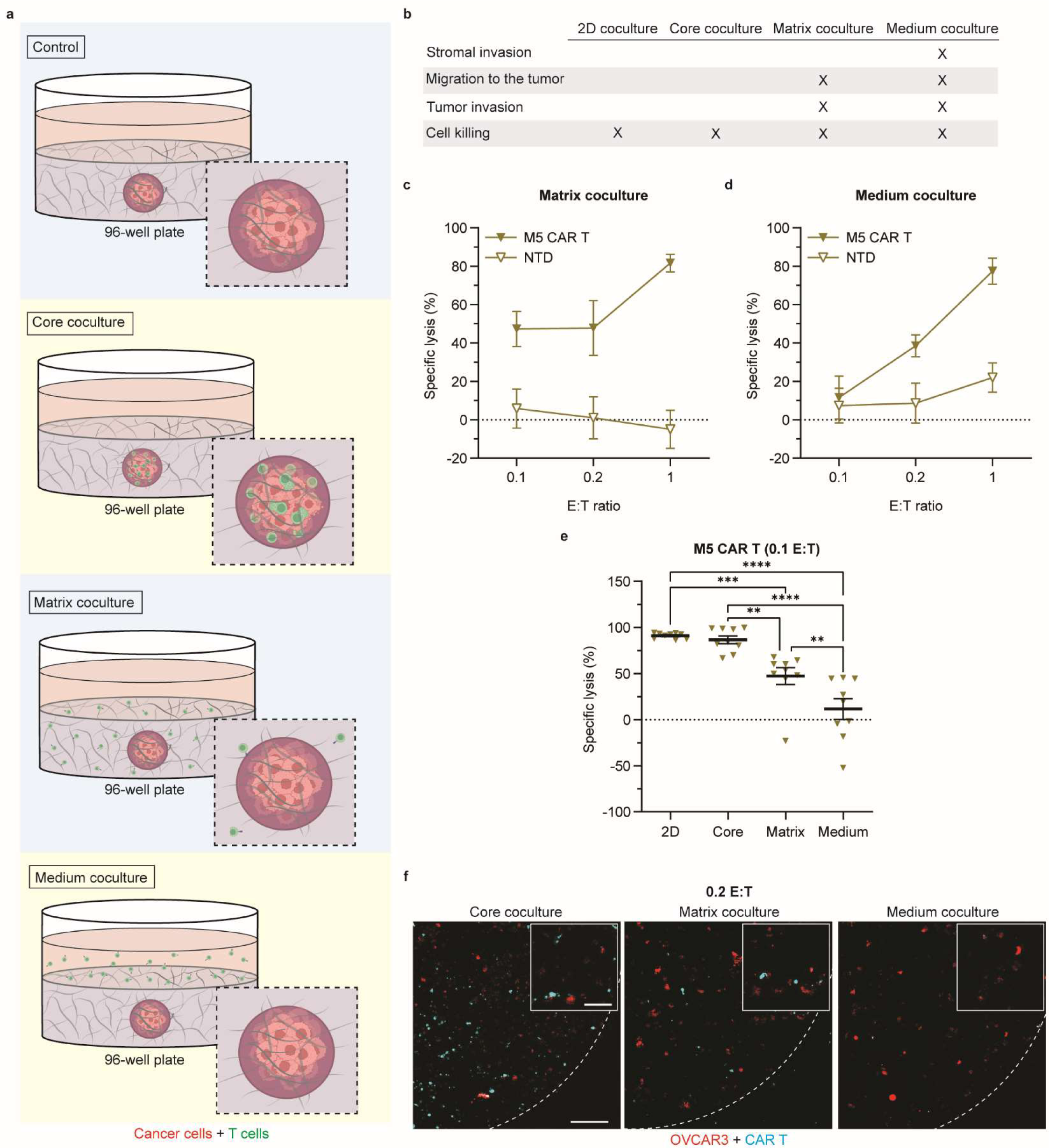
Multi-compartment cell infiltration/cytotoxicity assay. (a) Cartoons depicting the compartmentalization of the 3D infiltration/cytotoxicity assays. OVCAR3 ovarian cancer cells are grown in the Matrigel assembloid core and embedded in a bulk collagen I matrix. These cancer cells are luciferase-tagged so that their relative proliferation can be measured independent of immune cell proliferation. Three different coculture geometries simulate different steps of immune cell infiltration of a solid tumor: core coculture (second panel – T cells are seeded in the assembloid core), matrix coculture (third panel – T cells are seeded in the collagen matrix), and medium coculture (fourth panel – T cells are seeded in the medium on top of the assembloid). (b) Each cytotoxicity assay geometry simulates different steps of immune cell infiltration of a solid tumor. The cytotoxic effect of M5 CAR T and non-transduced (NTD) T cells against OVCAR3 ovarian cancer cells in the (c) matrix coculture, and (d) medium coculture on day 2. Specific lysis calculation is described in Methods. E:T, Effector:Target ratio. N = 3, n = 3. Data are mean ± SEM. (e) The cytotoxicity of M5 CAR T cells against OVCAR3 ovarian cancer cells in 2D and all 3D geometries. E:T ratio, 0.1. N = 3, n = 3. Data are mean ± SEM. Statistical test: two-way ANOVA with comparisons between all groups, ****P < 0.0001, ***P ≤ 0.001, **P ≤ 0.01. (f) Assembloids of each coculture geometry were imaged on day 2. M5 CAR T cells (cyan) are grown in coculture with OVCAR3 ovarian cancer cells (red) in the core coculture (left), matrix coculture (middle) and medium coculture (right) geometries. E:T ratio, 0.2. Dotted lines indicate the boundary of the assembloid cores. Images are maximum intensity projections of stacks of fluorescence confocal images. 10X magnification. Scale bar, 150 µm. Inset, 60 µm.

We first simulated T cells that had already migrated into the tumor and had come into direct contact with cancer cells to measure their 3D cytotoxicity (described above). In this core coculture geometry, T cells and OVCAR3 cancer cells were premixed and directly seeded in the core of the assembloid (Fig. 3a second panel – core coculture, Supplementary Fig. 2b). This model is a direct translation of the 2D standard cytotoxicity assay to a 3D system where immune cell-cancer cell interactions are immediately possible and 3D cell-ECM interactions are added, but the invasion and migration events required for immune infiltration of a solid tumor are bypassed. This geometry of the 3D cytotoxicity assay studied cell recognition and killing of cancer cells by immune cells. We observed more than 85% killing of OVCAR3 cells by CAR T cells at all E:T ratios (Fig. 2c).

We then simulated T cells that had arrived at the collagen-rich stroma near the tumor, but had not yet invaded the tumor. To achieve this, cancer cells were seeded alone in the assembloid core, while T cells were seeded in the surrounding collagen I matrix (Fig. 3a third panel – matrix coculture, Supplementary Fig. 2c). This geometry studies two immune infiltration events - migration to the tumor and tumor invasion - in addition to cytotoxicity (Fig. 3b). T cells migrate to the tumor assembloid core, invade the boundary of the core, and only then recognize and kill cancer cells. After two days, we observed 47-82% cell killing by CAR T cells (Fig. 3c) compared to > 85% when the cancer cells and CAR T cells were premixed (Fig. 2c), and the cytotoxicity of NTD T cells in this matrix coculture was negligible, confirming that CAR T cytotoxicity resulted from targeting of mesothelin expressing cells [9,22,36,37,44,45] (Fig. 3c). The reduced cytotoxicity of CAR T cells in matrix coculture from the cytotoxicity observed in core coculture could be caused by the time required for these immune cells to migrate to and then infiltrate the tumor assembloid core. In the core coculture (first geometry shown above), these processes were bypassed, enabling a more immediate cytotoxic effect.

The level of cytotoxicity also differed with the ratio of effector T cells to target cancer cells (E:T ratio). In the matrix coculture (Fig. 3c), this effect became evident. When seeded at lower E:T ratios, fewer immune cells were able to successfully navigate the ECMs to invade the tumor (Supplementary Fig. 1b), so the efficiency of the effector function was reduced.

Finally, we simulated T cells that have to undergo the entire infiltration process post-extravasation. The final design of the 3D infiltration/cytotoxicity assay seeded cancer cells in a Matrigel core and embedded the core in a bulk collagen matrix without immune cells. Immune cells were then deposited in culture medium on top of the assembloid (Fig. 3a fourth panel – medium coculture, Supplementary Fig. 2d). This geometry simulates stromal invasion, which immediately follows breaching of the endothelium in situ, in addition to the events captured in the matrix coculture system (Fig. 3b). We found that this geometry further reduced the cytotoxic impact of CAR T cells, with the lowest E:T ratio showing a cytotoxicity of < 20% after two days (Fig. 3d). The dose dependence of CAR T cell cytotoxicity was again reflected in this geometry as nearly 80% cell killing by CAR T cells was achieved at the highest E:T ratio.

These different geometries study multiple steps of the immune infiltration cascade and demonstrate how the assembloid architecture can be manipulated to isolate specific immune motility events or cell-ECM interactions. However, the addition of multiple steps in the immune infiltration cascade produces compounded delays in the ability of these cells to elicit a cytotoxic effect. The core coculture (first geometry) facilitates immediate cell killing. The matrix (second geometry) and medium coculture (third geometry) simulate additional invasion steps, so immune cell infiltration into the core and cytotoxicity decreased with each additional architectural complexity (Fig. 3e). The navigational barriers encountered in the later geometries reduced the number of immune cells that reached the cancer cells in the assembloid core by day 2 (Fig. 3f, Supplementary Fig. 1a). Immune cells in these geometries may need more time to successfully infiltrate the tumor, so we next explored the ability of our 3D assay to coculture cancer and immune cells for long time durations to better study the kinetics of cytotoxic activity.

### 3.3 The 3D infiltration/cytotoxicity assay allows for extended experimental durations

2D systems are limited by a maximum cell culture time that is dictated by when cells form a confluent monolayer, which typically occurs after 1-2 days. However, our 3D tumor assembloids can expand volumetrically [41] and are not bounded by the 2D area of the culture vessel. Cancer cells in the assembloid grow within the volumetric confinements of the Matrigel core, but can expand into the surrounding bulk collagen matrix if proliferation causes the total cell volume to exceed the core volume or cancer cells invade the collagen I matrix [41]. This mimics tumor expansion into the stroma in situ. Thus, our system permits long term (one week or longer [8,40–42]) culture that would not be otherwise possible in 2D systems which reach full confluency by day 2.

The assembloid geometries explored in our 3D assay demonstrated that introducing the ECM barriers experienced by cells in situ reduces tumor cell killing as the effector cells must cross each ECM barrier before eliciting an effector function (Fig. 3e). This difference in relative cell killing across geometries appeared with timescales held constant; however, these timescales can be extended to monitor longer-term immune interaction with the tumor microenvironment or observe immune cell kinetics under different stromal ECM properties (i.e., collagen density, gel thickness).

We hypothesized that cytotoxicity could be enhanced by increasing the duration of the 3D infiltration/cytotoxicity assay because the added time would be sufficient for effector cells to reach the tumor assembloid core and elicit their effector function. We tested two CAR T constructs, M5 [36,37,44,45] (used previously) and SS1 [37,52], which both target mesothelin expressing cells such as OVCAR3. When we cultured CAR T and OVCAR3 cells in the medium coculture geometry for six days, we observed a nearly 2.5-fold increase in M5 CAR T cytotoxicity compared to the results after two days of coculture and a more than four-fold increase in SS1 CAR T cytotoxicity (Fig. 4a). Hypoxia and nutrient competition are often encountered in the tumor microenvironment; although, we do not anticipate compounding effects from these factors as culture medium was replenished every 2 days. These conditions could be introduced in future studies using this model, however. Extending the experimental duration of these assembloid systems also means CAR T cell counts per condition can be reduced. Standard in vitro cytotoxicity assays attempt to saturate target cells with high E:T ratios (up to 100 E:T) to achieve quick cytotoxic effects in 2D [9]. We seeded 1 effector cell to every 10 target cells (0.1 E:T), which is lower than most 2D assays, and still observed a cytotoxic effect. Reducing the number of immune cells per condition allows for the study of additional TME conditions or immune cell-TME interactions within the same experiment or PBMC (peripheral blood mononuclear cell) donor.

**Figure 4.**
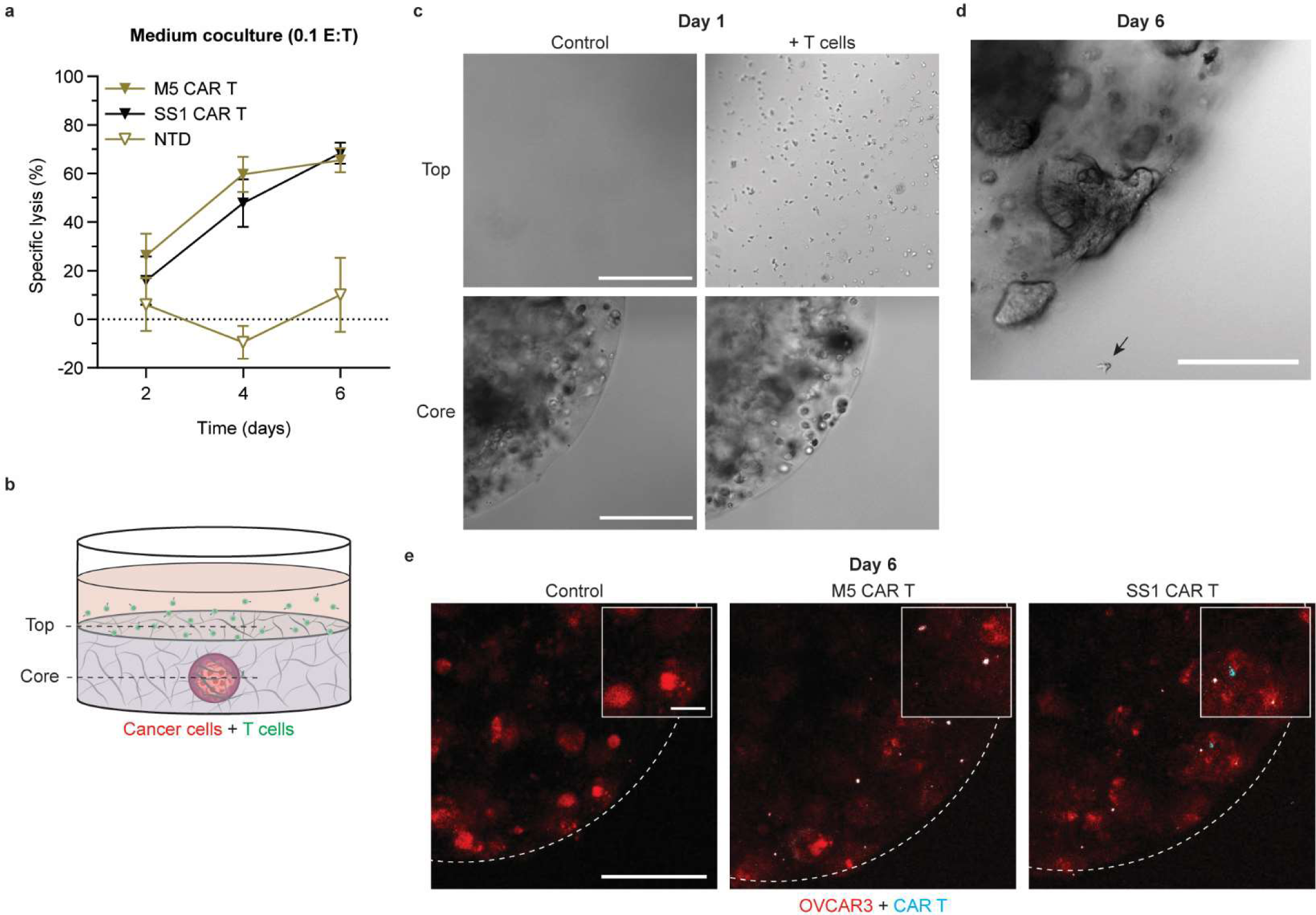
Extending experimental timelines enhances immune infiltration and cell killing. (a) The cytotoxicity of M5 CAR T cells against OVCAR3 ovarian cancer cells up to 6 days of medium coculture. E:T ratio, 0.1. N = 3, n = 3. Data are mean ± SEM. The luciferase activity for distinct samples was measured on days 2, 4, and 6. (b) Cartoon depicting immune cell addition in the medium on top of the assembloid bulk matrix and the assembloid core. (c) Effector cells were added to the top of the matrix on day 1 of coculture where they were not immediately in contact with the target cells in the assembloid core. Images are DIC microscopy. CAR T and NTD T cells were grown in medium coculture with OVCAR3 cells. 10X magnification. Scale bar, 250 µm. By day 6, effector cells d, reached and e, invaded the assembloid core. E:T ratio, 0.1. (d) 10X magnification, DIC. Scale bar, 150 µm. (e) Dotted lines indicate the boundary of the assembloid cores. OVCAR3 cells (red) and CAR T cells (cyan) in matrix coculture. Images are maximum intensity projections of stacks of fluorescence confocal images. 10X magnification. Scale bar, 250 µm. Inset, 50 µm.

On the first day of coculture, effector cells added in the assembloid medium were separated from cancer cells by a bulk collagen matrix (Fig. 4b,c). By day six, T cells had invaded the collagen matrix and could be found in the same z-plane as the assembloid core (Fig. 4d). We confirmed the infiltration of CAR T cells into the tumor assembloid core by confocal microscopy (Fig. 4e, Supplementary Fig. 1c). These visual confirmations were consistent with the cytotoxicity results and allowed for visual observation of immune infiltration events. Immune cell infiltration was enhanced by extending the timeline of matrix coculture, and effector-target cell contact was confirmed through microscopy images and luciferase activity.

### 3.4 Testing different cell types

Our analysis has focused on T cells and CAR T cells, including the M5 and SS1 benchmark CAR T constructs that target mesothelin-expressing cancer cells (i.e., OVCAR3). However, the versatility of the 3D assay also extends to shedding new light on the behavior of other immune or TME cell types and the mechanisms causing the success or failure of immune cells to target cancer cells in the solid TME [53].

The transition from 2D coculture to 3D core coculture doubled the cytotoxicity of naïve M0 macrophages (Fig. 5a). However, cancer-cell lysis by macrophages was limited by transitioning to geometries with additional hurdles (matrix and medium cocultures). Thus, our model reveals that 3D ECM components such as the basement membrane within the assembloid core can alter the immune response compared to 2D systems. The motility of M0 macrophages was highly limited (Supplementary Video 1), so these effector cells did not move from the compartment where they were seeded (Fig. 5b, Supplementary Fig. 3a). Even though M0 macrophages in the matrix and medium cocultures were stimulated by a 3D microenvironment, they were slow to invade the assembloid core. After three days, some M0 macrophages could be found invading the assembloid core (Fig. 5c, Supplementary Fig. 3b). Some M0 macrophages were able to invade the assembloid core when seeded in the assembloid matrix, which demonstrated that even immune cells with limited motility were able to infiltrate the assembloid tumor core where close cell-cell contact is possible and immune synapses can form. The low infiltration of M0 macrophages into the assembloid core also aligns with the biological role of macrophages. Most often, macrophages differentiate from monocytes locally instead of circulating or migrating as macrophages [1], so the limited motility of macrophages observed in our 3D assay matches their physiological role in cancer.

**Figure 5.**
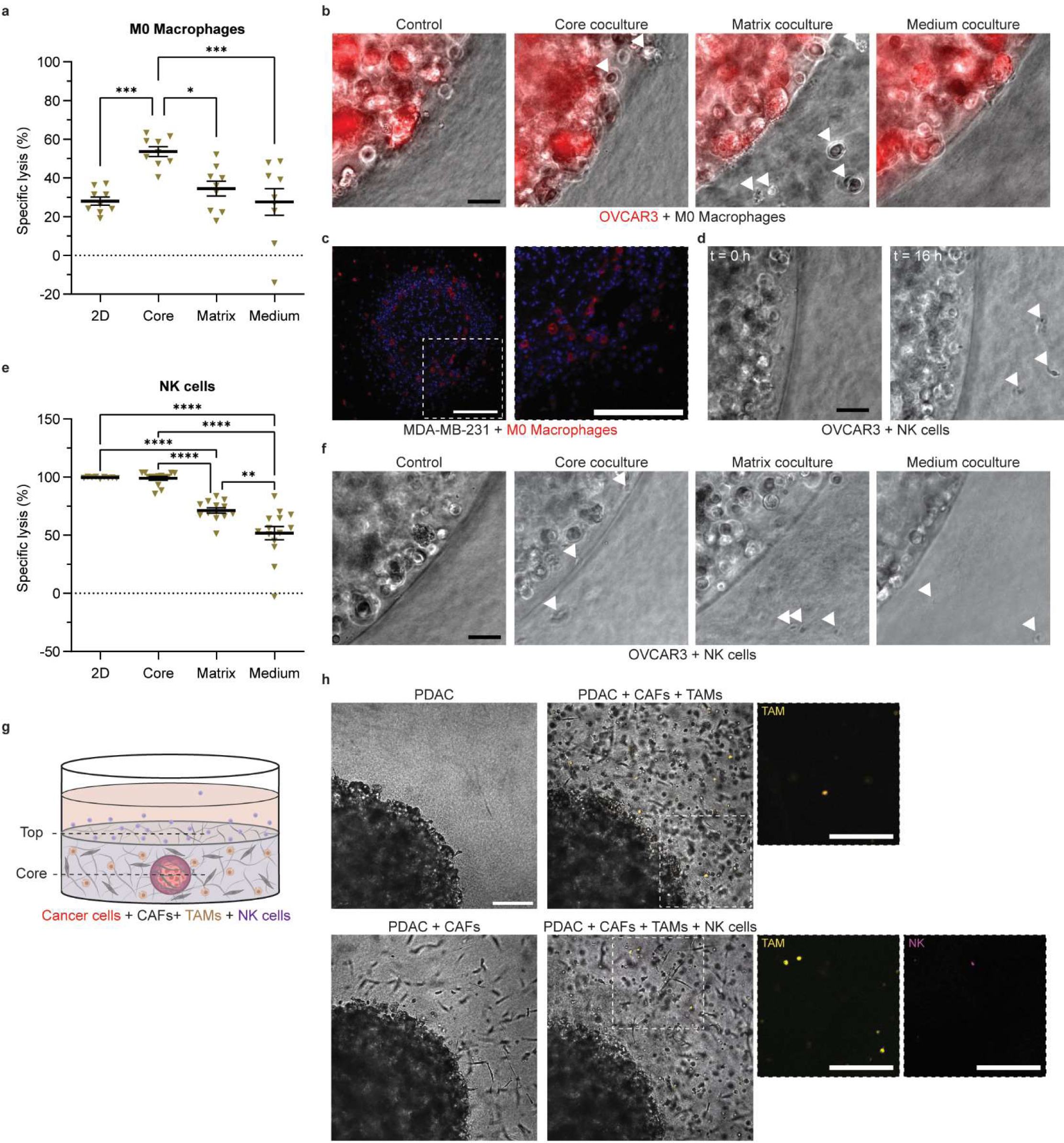
Cell killing efficiency differs by immune cell subtypes. (a) The cytotoxic effect of M0 Macrophages against OVCAR3 ovarian cancer cells in 2D and all 3D coculture geometries on day 2. E:T ratio, 1. N = 3, n = 3+, donors = 3. Data are mean ± SEM. Statistical test: two-way ANOVA with comparisons between all groups, ***P ≤ 0.001, *P ≤ 0.05. (b) M0 macrophages did not cross ECM boundaries by day 2. Images are phase contrast images overlayed with epifluorescence images showing OVCAR3 ovarian cancer cells in red. Arrowheads point to M0 macrophages. 10X magnification, Scale bar, 50 µm. (c) On day 3, M0 macrophages that were seeded in the matrix coculture geometry were found invading the assembloid core. Images are fluorescence images of FFPE sections of assembloids where nuclei are stained in blue and M0 macrophages are stained in red. The cancer cells in the assembloid core are MDA-MB-231 breast cancer cells. Scale bars, 200 µm. (d) NK cells were motile within the multi-compartment assembloid. NK cells were not visible near the boundary of the OVCAR3 core at time 0 h, but appear by 16 h. Images are phase contrast images. Arrowheads point to NK cells. 10X magnification, Scale bar, 50 µm. (e) The cytotoxic effect of NK cells against OVCAR3 ovarian cancer cells in 2D and all 3D coculture geometries on day 2. E:T ratio, 1. N = 3, n = 3+, donors = 2. Data are mean ± SEM. Statistical test: two-way ANOVA with comparisons between all groups, ****P < 0.0001, **P ≤ 0.01. (f) NK cells crossed ECM boundaries by day 2. Images are phase contrast images. Arrowheads point to NK cells. 10X magnification, Scale bar, 50 µm. (g) Cartoon depicting an assembloid with PDAC cells in the core, CAFs and TAMs seeded in the bulk matrix, and NK cells seeded in medium on top of the bulk matrix. (h) DIC images on day 4 of assembloids with PDAC cells in the assembloid core, CAFs and TAMs (yellow) seeded in the bulk matrix, and NK cells (magenta) seeded in the medium on top of the bulk matrix. 10X magnification, DIC with fluorescence microscopy overlay. All scale bars, 250 µm.

In contrast to macrophages, natural killer (NK) cells moved quickly throughout the collagen matrix (Fig. 5d) and readily crossed the Matrigel-collagen boundary (Supplementary Video 2). The cytotoxicity of NK cells in the 3D core coculture was equivalent to their cytotoxicity in 2D coculture (Fig. 5e). NK cells in matrix coculture and medium coculture also elicited a cytotoxic response toward OVCAR3 cancer cells and could be found at the boundary of the assembloid core within two days of coculture (Fig. 5f, Supplementary Fig. 3c); although, this effect was attenuated compared to the core coculture. This was consistent with the time delay in cytotoxicity observed in CAR T cell assays (Fig. 4a) and reinforced the importance of assessing invasion and migration through the ECM when evaluating the role of immune cells in the TME.

Our 3D cytotoxicity assay can also be used to study immune-TME interactions in less explored subtypes of cancer. Gestational trophoblastic neoplasia (GTN) is one such type of gynecologic cancer where the tumor and surrounding stroma is infiltrated by many different immune populations [54], and thus, could be amenable to immunotherapies [54,55]. We assessed the cytotoxicity of macrophages (Supplementary Fig. 4a) and NK cells (Supplementary Fig. 4b) in coculture with GTN cells. M1 macrophages [56,57] and NK cells elicited an expected cytotoxic effect in core coculture; however, their cytotoxicity in other coculture geometries was negligible, which differed from the cytotoxicity patterns observed by NK cells against OVCAR3 ovarian cancer cells. Thus, future studies should investigate the final steps of infiltration of these immune subtypes into the GTN tumor.

Other experimental parameters of the 3D combined infiltration/cytotoxicity assay are also tunable. Above, we have demonstrated the versatility of the assembloid model through modifications that facilitated the study of different immune infiltration events via different assembloid geometries (Fig. 3, Supplementary Fig. 1a), E:T ratios (Fig. 2c,f; Fig. 3c-d, Supplementary Fig. 1b), experimental time durations (Fig. 4), immune cell populations (Fig. 5a-f), and cancer types (Supplementary Fig. 3,4). The assay can be further tuned to reflect the cellular heterogeneity of the tumor microenvironment and disease-related changes to the ECM [58]. Additional stromal cells can be incorporated, or multiple immune cell types can be simultaneously cocultured with cancer cells. To demonstrate the cellular heterogeneity that can be achieved in this assay, we cocultured pancreatic ductal adenocarcinoma cancer-associated fibroblasts (PDAC CAFs) in the collagen matrix surrounding a PANC-1 PDAC assembloid core. Tumor-associated macrophages (TAMs) (CD206+, CD163+ and CD80- [59–61], Supplementary Fig. 5a-d) and NK cells were physiologically incorporated into the bulk matrix and culture medium, respectively, to simulate an example microenvironment with complex multi-cellular interactions (Fig. 5g,h). In this coculture system, CAFs and TAMs remained isolated from PDAC cells in the adjacent collagen compartment, and NK cells deposited in the medium navigated through the cellular and ECM stroma to reach the assembloid core. Interactions between CAFs and immune cells in the tumor microenvironment have been shown to contribute to tumor progression [21] and can be systematically analyzed in future studies using this 3D model.

## 4. Discussion

Our 3D in vitro infiltration/cytotoxicity assay recapitulates several steps involved in immune cell infiltration of the tumor, which results in more realistic cytotoxicity toward cancer cells than traditional 2D cell killing assays. The 3D assembloid model used in the assay facilitates the study of effector cell-TME interactions and their role in immune infiltration/cytotoxicity in solid tumors. We demonstrate how the model can be tuned to simulate different stages of immune cell infiltration, individually and as an infiltration cascade. The standard of the field measures the cytotoxicity of forced effector-target cell interactions in an artificial 2D microenvironment [9–11] without considering these infiltration steps and 3D cell-ECM interactions. One predominant method to address the gap between cell culture plastic or glass (2D) and the ECM found in solid tumors is to provide a coating of relevant ECM prior to cell seeding to better recapitulate the ECM sensed by cells in vivo [62]. However, this approach is limited to bidirectional cell-ECM stimulation and lacks migration/invasion requirements. Additionally, the stiffness sensed by cancer cells is the same as plastic or glass (∼GPa) [63], leading to different behavior than what would be observed with stiffnesses of physiological relevance [16]. One possible solution has been to plate cells on gels of desired stiffness, which can also be given a coating of desired ECM to further increase their physiological relevance [64]. Still, these systems are limited to 2D cell-ECM interactions and neglect the critical infiltration and migration steps immune cells must overcome to reach cancer cells. These points and other advantages/disadvantages of the standard 2D assay and our 3D assay are outlined in Table 1.

**Table 1.**
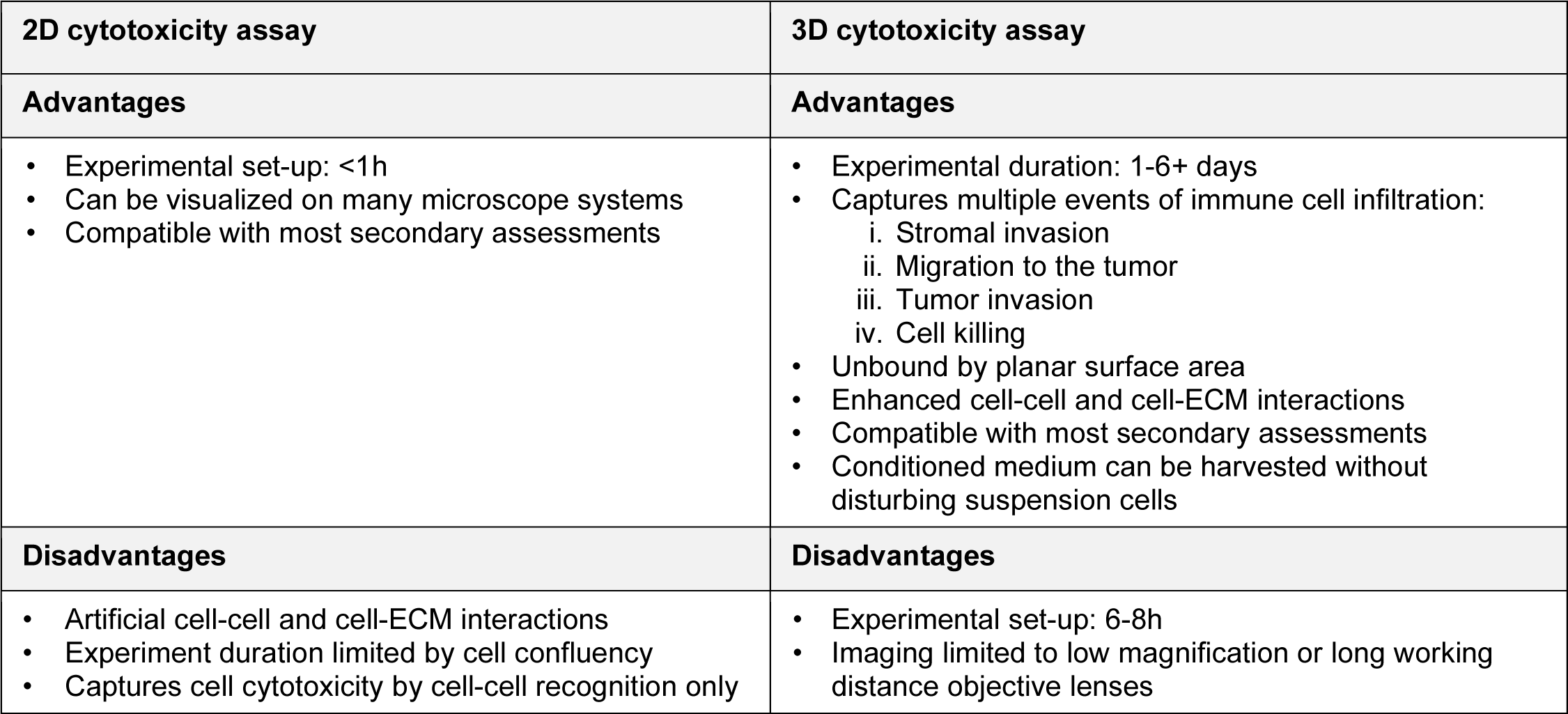
Advantages and disadvantages of the 2D and 3D assays.

| 2D cytotoxicity assay | 3D cytotoxicity assay |
| --- | --- |
| **Advantages** | **Advantages** |
| • Experimental set-up: <1h<br>• Can be visualized on many microscope systems<br>• Compatible with most secondary assessments | • Experimental duration: 1-6+ days<br>• Captures multiple events of immune cell infiltration:<br>i. Stromal invasion<br>ii. Migration to the tumor<br>iii. Tumor invasion<br>iv. Cell killing<br>• Unbound by planar surface area<br>• Enhanced cell-cell and cell-ECM interactions<br>• Compatible with most secondary assessments<br>• Conditioned medium can be harvested without disturbing suspension cells |
| **Disadvantages** | **Disadvantages** |
| • Artificial cell-cell and cell-ECM interactions<br>• Experiment duration limited by cell confluency<br>• Captures cell cytotoxicity by cell-cell recognition only | • Experimental set-up: 6-8h<br>• Imaging limited to low magnification or long working distance objective lenses |

Our 3D assay provides additional insights into the physiological interactions between immune cells and the TME. While the cytotoxicity of immune cells when in close proximity to cancer cells is an important feature of immune interactions with the tumor, we must also consider that these cells need to infiltrate the tumor before such killing can occur [28]. Infiltration is one of the leading current challenges in cell immunotherapies limiting their success against solid tumors [4,14,30]. Infiltration alone can significantly enhance the efficacy of CAR T cells against solid tumors, as demonstrated in a recent work by some of the authors [4]. Yet, infiltration is not addressed in the standard 2D assay or other recently established 3D cytotoxicity assays [22–24]. Our medium coculture assay allows for simulation of these key infiltration events. The spatial distribution of in situ immune and cancer cells, which can be observed from tissue sections (immune cells localized within the tumor vs. surrounding the tumor), is important when studying the molecular and physical mechanisms of immune cell cytotoxicity and can be simulated using the assembloid geometries we demonstrated here. This versatile 3D cytotoxicity assay offers a platform where the steps of immune infiltration can be independently measured via endpoint cell viability assays and studied in real-time using many microscopy techniques.

The 3D system could also be utilized to uncover new biology pertaining to the migration of immune cells, their invasion through ECM, and their interactions with cancer cells. In our analysis, the effector response to target cells varied by effector cell subtype and could be changed through the introduction of specific 3D cell-ECM interactions. This varied by cancer type, ECM composition, and the distance between immune cells and cancer cells. Macrophages, for example, were less cytotoxic toward OVCAR3 ovarian cancer cells, in part because they struggled to infiltrate the TME, but could be stimulated by a 3D basement membrane-based microenvironment. CAR T cells and NK cells were highly cytotoxic toward OVCAR3 ovarian cancer cells and successfully infiltrated the tumor assembloid to perform their effector functions, but NK cells were less cytotoxic toward JAR GTN cells. The ECM composition was unchanged in our GTN and ovarian cancer assays, so this identifies an opportunity for such 3D in vitro systems to explore the mechanical cancer cell-NK cell interactions and soluble factor crosstalk between effector and target cells that are different for JAR (GTN) and OVCAR3 (ovarian cancer) cells. The exact mechanisms of NK cell killing in this context and how these mechanisms differ between OVCAR3 and JAR cells should be further explored in future studies applying this model. This new 3D assay is a powerful tool for customizable and informative study of the biology of immune cells in the TME and their infiltration/cytotoxicity in solid tumors.

Using high throughput 3D spatial mapping of whole organs and assembloids, we have previously demonstrated that the multi-compartment assembloid method more accurately recapitulates the compartmentalization, cell and ECM composition, and complex morphology of tissues than other cell culture methods [40]. Thus, the similar assembloid model used in this 3D combined infiltration/cytotoxicity assay can be intricately tuned to reflect the tissue anatomy and customized to expand mechanistic understanding of different cell-ECM interactions on cell motility that have not yet been explored in the context of immunology [27,29]. This includes introducing several components of the tumor microenvironment that have been shown to modulate the immune response, including fibroblasts and pro-tumorigenic immune cells (i.e., TAMs) [65]. These modifications were introduced in this work, but should be mechanistically explored in greater detail in future studies. Our model also permits spatial control over the introduction of these different cell types by seeding them in outer compartments as opposed to the assembloid core with cancer cells. By matching the in situ compartmentalization of cell types and ECM [15], we can model immune subtype-specific patterns of TME infiltration [53]. Extravasation of immune cells from a blood vessel into the tumor could also be incorporated into future versions of the assay by introducing a monolayer of endothelial cells and/or smooth muscle cells on top of the collagen matrix prior to seeding immune cells in the medium coculture geometry. We demonstrated some applications of this assay when studying the infiltration of different immune cell types in the ovarian cancer TME; however, this assay could also be used to explore the roles of ECM composition or immune cell-driven remodeling of the ECM [27,66,67].

## 5. Conclusion

Here, we focus our introduction of the 3D combined infiltration/cytotoxicity assay on cell-ECM interactions and cell motility, but its potential applications expand into many areas of cancer immunology. Nutrient competition as a result of immune infiltration of a solid tumor and interactions with resident stromal cells (i.e., CAFs) are broad examples of applications which were not explored here but could be explored through future customized versions of this model. This model could also be used to study molecular or antibody-based targets in the TME, such as immune checkpoint inhibitors (i.e., anti-PD1 or anti-PD-L1) [68], inflammatory cytokines [4], and other drug screens [42]. While the data presented here studies immune infiltration and cytotoxicity against immortalized cancer cell lines, the assembloid model supports primary cancer cell culture [8] and can also be used in future studies of inter-patient heterogeneity of immune response via personalized medicine using patient-derived cancer cells and PBMCs.

Such assays studying the infiltration of immune cells are necessary for advancing engineered cell therapies, particularly their success against solid tumors in the clinic. Recent work has shown that CAR T cells that infiltrate solid tumors more successfully kill cancer cells than CAR T cells that only reach the tumor periphery, despite equivalent cytotoxicity in 2D cell recognition assays [4]. Thus, 3D assays that address cell infiltration, such as the 3D infiltration/cytotoxicity assay described here, are critically important.

## CRediT authorship contribution statement

**Ashleigh J. Crawford:** Conceptualization, Formal analysis, Investigation, Methodology, Project administration, Supervision, Validation, Visualization, Writing – original draft, Writing – review & editing. **Adrian Johnston:** Conceptualization, Formal analysis, Investigation, Methodology, Supervision, Validation, Visualization, Writing – review & editing. **Wenxuan Du:** Formal analysis, Investigation, Supervision, Validation, Visualization, Writing – review & editing. **Eban A. Hanna:** Formal analysis, Investigation, Validation, Visualization, Writing – review & editing. **David Schell:** Formal analysis, Investigation, Validation, Visualization, Writing – review & editing. **Zeqi Wan:** Investigation. **Ting-Hsi Chen:** Investigation. **Fan Wu:** Investigation, Writing – review & editing. **Kehan Ren:** Investigation, Visualization, Writing – review & editing. **Yeongseo Lim:** Investigation. **Praful Nair:** Supervision, Visualization, Writing – review & editing. **Denis Wirtz:** Conceptualization, Funding acquisition, Methodology, Project administration, Supervision, Visualization, Writing – original draft, Writing – review & editing.

## Acknowledgements

The authors thank J. Phillip, B. Vogelstein, and T. Nichakawade for their important feedback. We also thank all members of the Phillip lab who generously allowed use of microscopes throughout this work. The authors acknowledge the following funding sources: the National Institute of Health (R01CA174388) to D.W., the National Cancer Institute (U54CA143868 and U54CA268083) to D.W., the National Institute of Arthritis and Musculoskeletal and Skin Diseases (U54AR081774) to D.W., and the National Institute on Aging (U01AG060903) to D.W. The authors also acknowledge equipment support from the Integrated Imaging Center at Johns Hopkins University.

## Data availability statement

All data are available in the main text or supporting materials. All raw data and cell lines are available upon request.

