## Supplementary Information for "A 3D in vitro assay to study combined immune cell infiltration and cytotoxicity"

**Corresponding author*

[+]*Indicates authors contributed equally*

### Supplementary Information

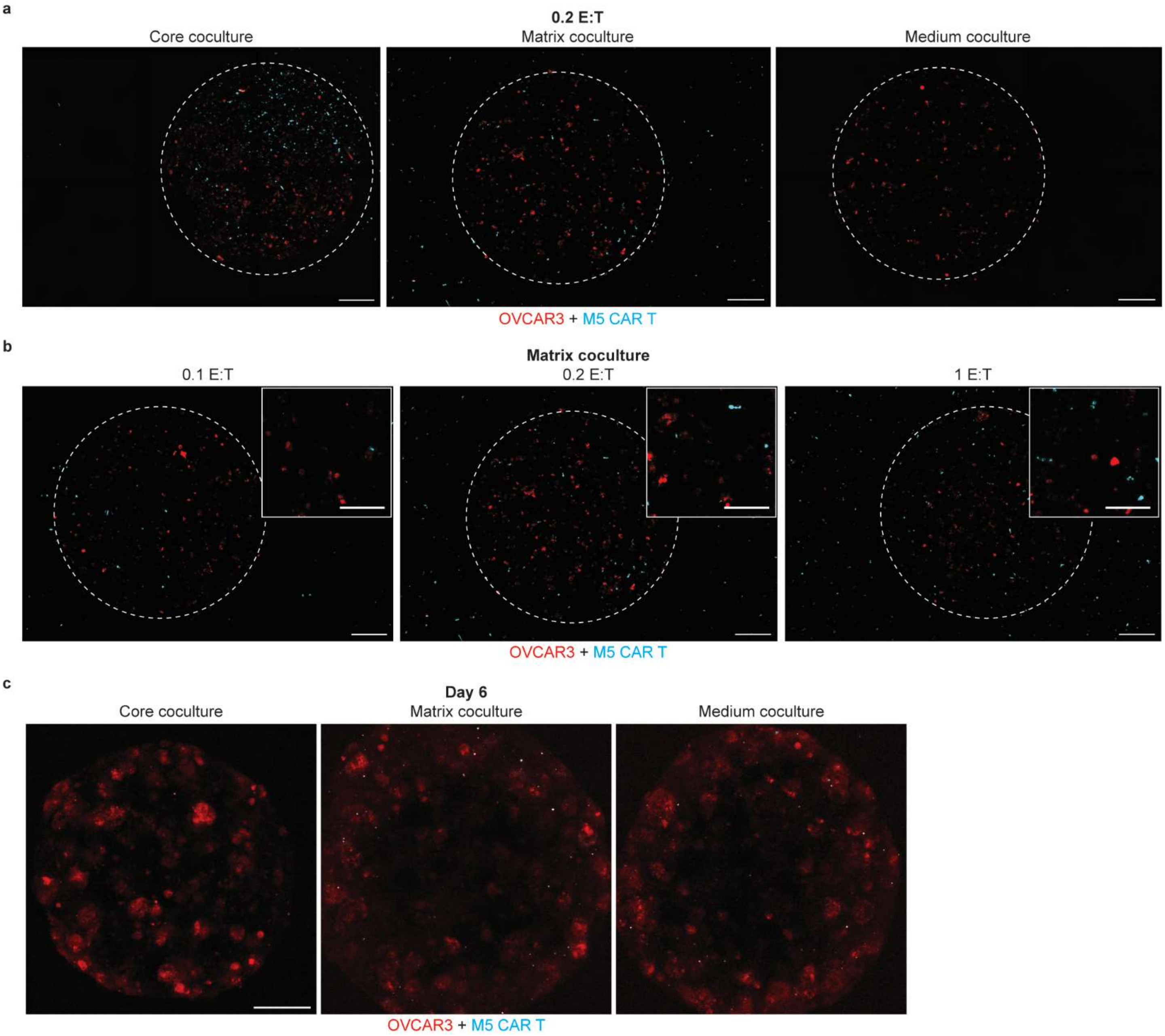


**Supplementary Figure 1. 3D cytotoxicity assay.** (a) Assembloids of each coculture geometry on day 2. M5 CAR T cells (cyan) are grown in coculture with OVCAR3 ovarian cancer cells (red) in the core coculture (left), matrix coculture (middle) and medium coculture (right) geometries. Dotted lines indicate the boundary of the assembloid cores. E:T ratio, 0.2. Images are maximum intensity projections of stacks of fluorescence confocal images. 10X magnification. Scale bars, 250 μm. (b) Matrix coculture assembloids at different E:T ratios on day 2. M5 CAR T cells (cyan) are grown in coculture with OVCAR3 ovarian cancer cells (red). Dotted lines indicate the boundary of the assembloid cores. Images are maximum intensity projections of stacks of fluorescence confocal images. 10X magnification. Scale bars, 250 μm. Inset, 125 μm. (c) Day 6 medium coculture assembloids where effector cells (cyan) have infiltrated the tumor assembloid and are in contact with or close proximity to OVCAR3 cancer cells (red). Images are maximum intensity projections of stacks of fluorescence confocal images. 10X magnification. Scale bar, 250 μm.

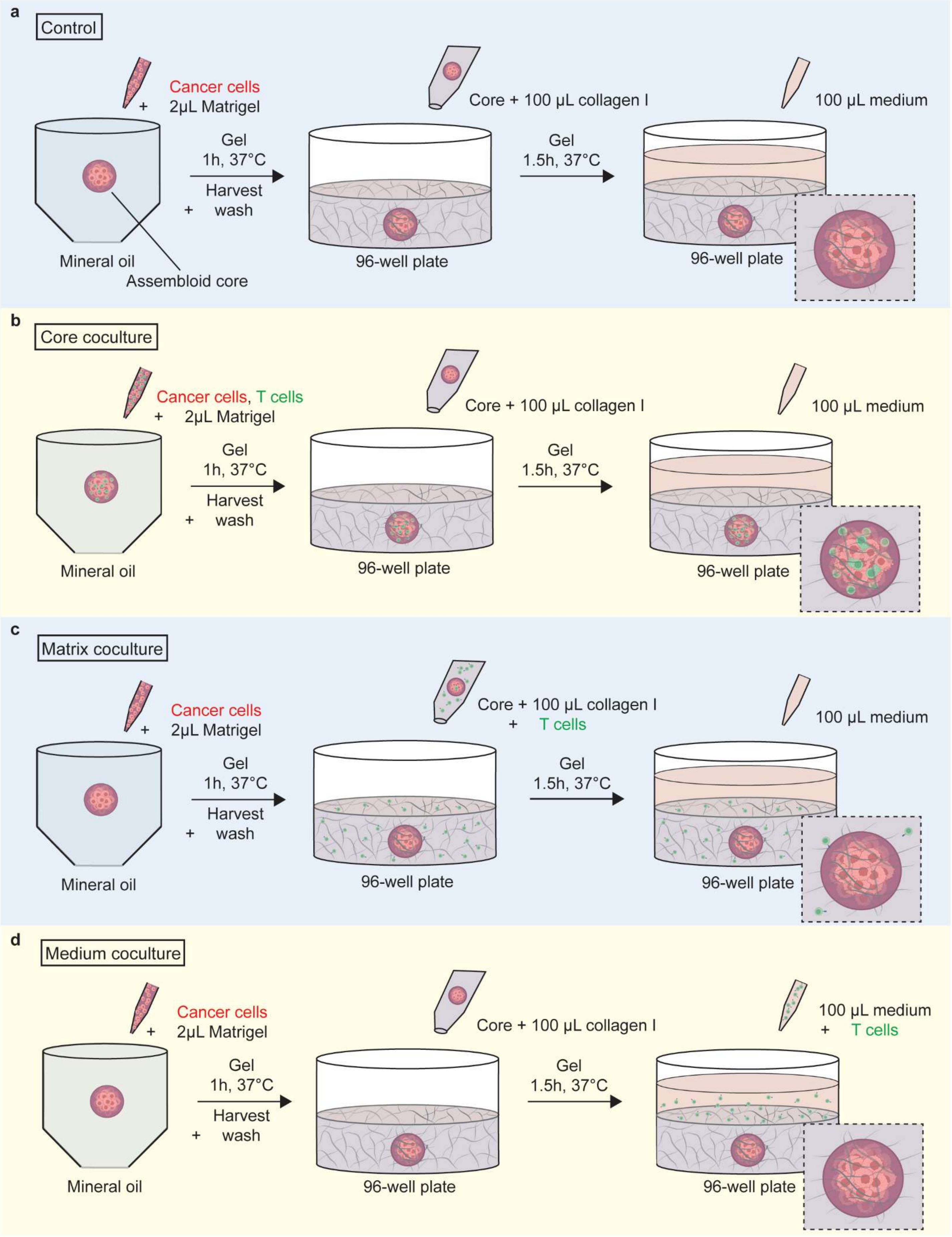
a
Control
Cancer cells
+ 2μL Matrigel
Gel
1h, 37°C
Harvest
+ wash
Mineral oil
Assembloid core
Core + 100 μL collagen I
96-well plate
Gel
1.5h, 37°C
100 μL medium
96-well plate
b
Core coculture
Cancer cells, T cells
+ 2μL Matrigel
Gel
1h, 37°C
Harvest
+ wash
Mineral oil
Core + 100 μL collagen I
96-well plate
Gel
1.5h, 37°C
100 μL medium
96-well plate
c
Matrix coculture
Cancer cells
+ 2μL Matrigel
Gel
1h, 37°C
Harvest
+ wash
Mineral oil
Core + 100 μL collagen I
+ T cells
96-well plate
Gel
1.5h, 37°C
100 μL medium
96-well plate
d
Medium coculture
Cancer cells
+ 2μL Matrigel
Gel
1h, 37°C
Harvest
+ wash
Mineral oil
Core + 100 μL collagen I
96-well plate
Gel
1.5h, 37°C
100 μL medium
+ T cells
96-well plate

**Supplementary Figure 2. Techniques for generating multi-compartment assembloid geometries.** Cartoons depicting the techniques used to generate multi-compartment assembloids for each geometry. (a) Control assembloids contain target cells in a 2 µL Matrigel core which is embedded in a 100 µL collagen I matrix in a 96 well plate. No effector cells are included in the control condition, which is used as the live and dead (treated with TritonX) controls. Cancer cells are luciferase-tagged in all conditions so that cancer cell proliferation can be measured independent from immune cell proliferation. (b) Core coculture assembloids mix effector cells with target cells in the assembloid core. (c) Matrix coculture contains only target cells in the assembloid core, but mixes effector cells into the bulk collagen matrix. (d) Medium coculture is assembled with no effector cells in the core or bulk matrix, but effector cells are added in the medium on top of the assembloid.

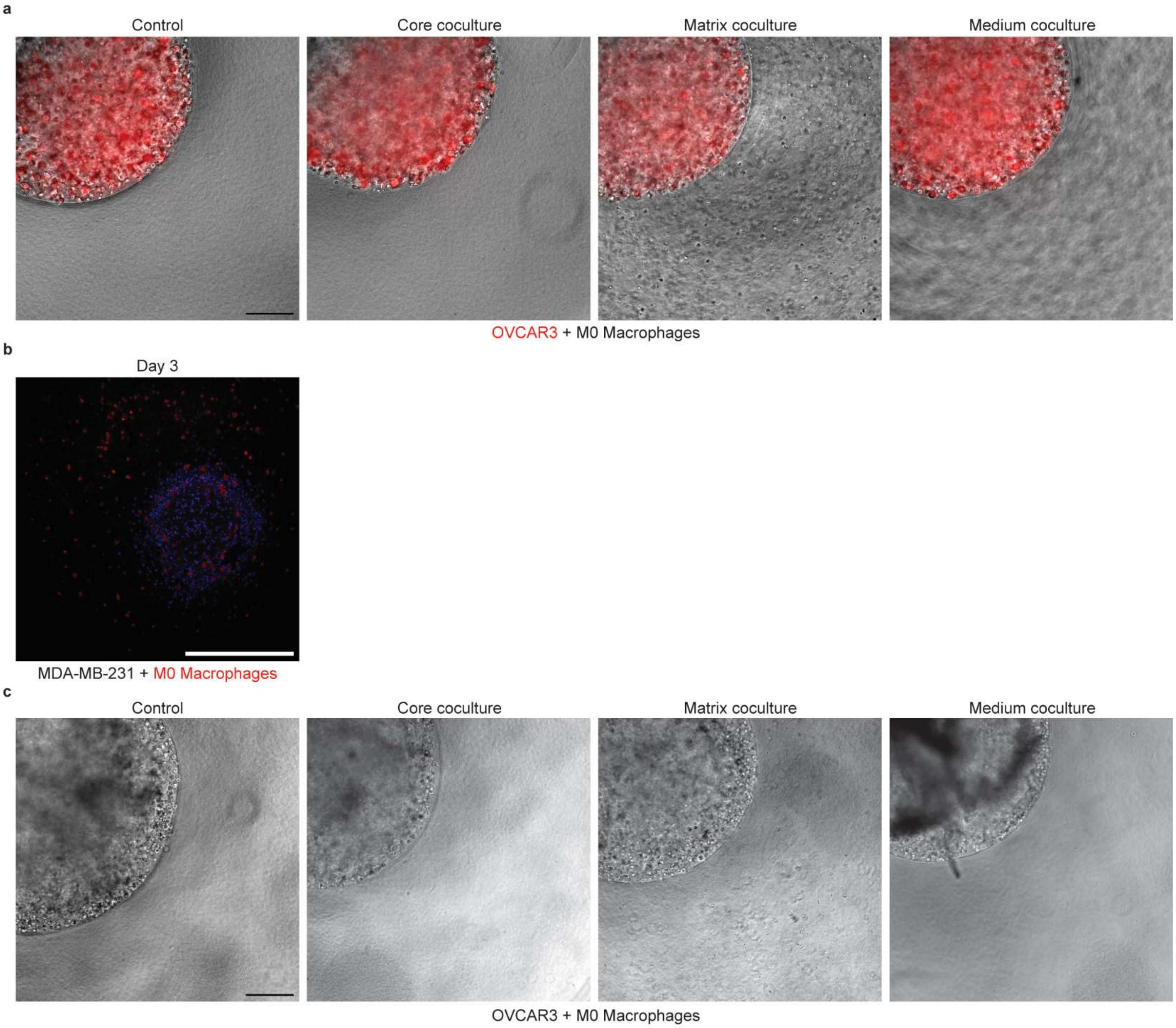


**Supplementary Figure 3. 3D cytotoxicity assay with macrophages and NK cells.** (a) M0 macrophages remain mostly in the assembloid region where they were seeded by day 2. Images are phase contrast images overlayed with epifluorescence images showing OVCAR3 cells in red. 10X magnification, Scale bar, 300 μm. (b) By day 3, M0 macrophages seeded in the matrix coculture geometry began to invade the assembloid core and were found at the periphery. Images are fluorescence images of FFPE sections of assembloids where nuclei are stained in blue and M0 macrophages are stained in red. The cancer cells in the assembloid core are MDA-MB-231 breast cancer cells. Scale bar, 500 μm. (c) NK cells move throughout the assembloid and cross ECM barriers by day 2. Images are phase contrast images. 10X magnification, Scale bar, 300 μm.

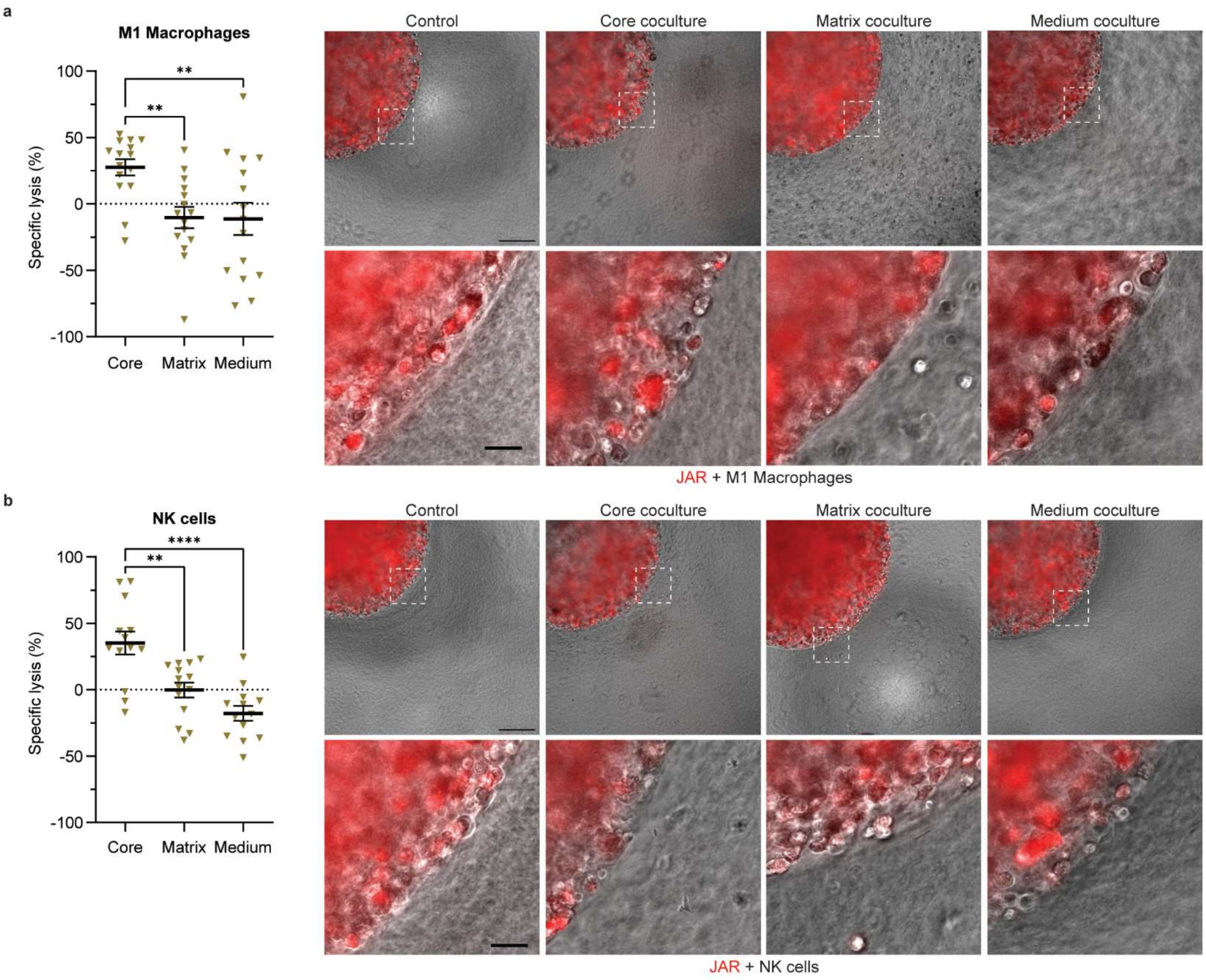


**Supplementary Figure 4. The 3D assay screens immune infiltration and cytotoxicity toward gestational trophoblastic neoplasia.** (a) M1 macrophages have a limited cytotoxic effect against JAR cancer cells (GTN) and stay within the assembloid compartment they were seeded in. Specific lysis calculation is described in Methods. E:T ratio, 1. N = 3, n = 3+, donors = 3. Data are mean ± SEM. Statistical test: two-way ANOVA with comparisons between all groups, **P ≤ 0.01. Images are day 2 phase contrast images overlayed with epifluorescence images showing JAR cells in red. 10X magnification, Scale bar, 300 µm and 50 µm (inset). (b) NK cells elicit a cytotoxic effect against JAR cells in 3D core coculture and move throughout assembloid compartments. Specific lysis calculation is described in Methods. E:T ratio, 1. N = 3, n = 3+, donors = 3. Data are mean ± SEM. Statistical test: two-way ANOVA with comparisons between all groups, ****P < 0.0001, **P ≤ 0.01. Images are day 2 phase contrast images. 10X magnification, Scale bar, 300 µm and 50 µm (inset).

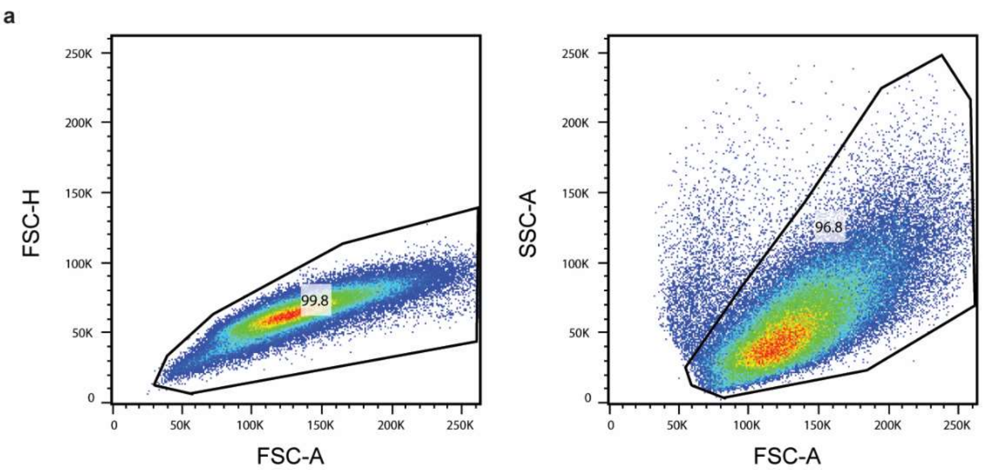


d

Positive cells (%)

| | TAM | M2 | Isotype control |
|---|---|---|---|
| **CD206** | 79.9 | 76.4 | 0.16 |
| **CD163** | 53.2 | 57.0 | 0.11 |
| **CD80** | 0.64 | 0.61 | 0.32 |

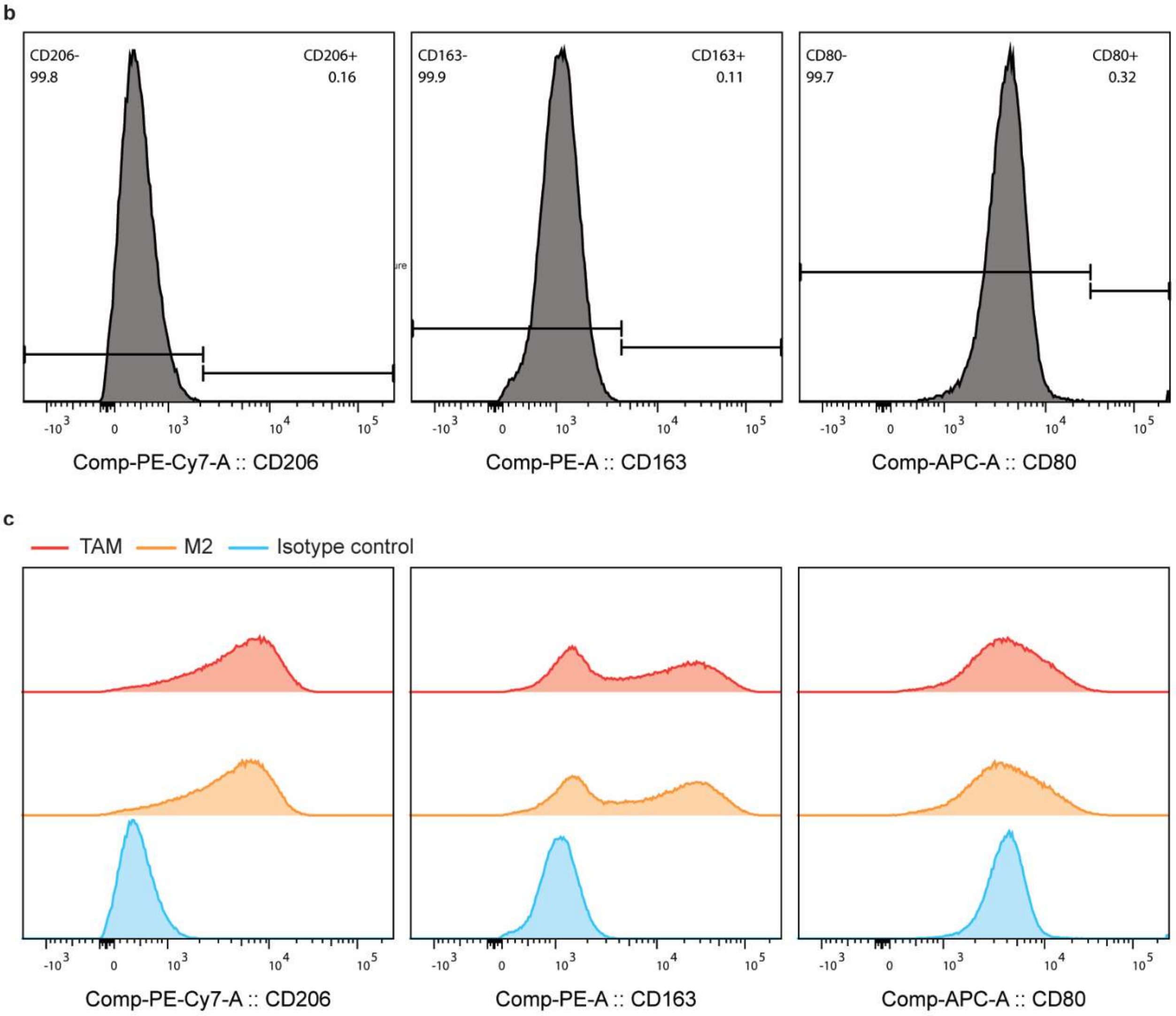


**Supplementary Figure 5. Tumor associated macrophages are confirmed by flow cytometry.** (a) Gating strategy for isotype control single cells (left) and live cells (right). (b) The boundaries for CD206, CD163, and CD80 (from left to right) were determined by the isotype control. (c) Histograms of select macrophage markers. From left to right, CD206, CD163, and CD80. TAMs and M2 macrophages are CD206+, CD163+ and CD80- [1–3]. (d) Percentage of TAM, M2 and isotype control cell populations that were positive for CD206, CD163, and CD80 after gating by the isotype control.

**Supplementary Video 1.** Primary human macrophages (M0) in the collagen compartment of a matrix coculture assembloid. OVCAR3 ovarian cancer cells (red) were seeded in the assembloid core with M0 macrophages in the collagen matrix. Scale bar, 500 μm.

**Supplementary Video 2.** Primary human NK cells in a matrix coculture assembloid. OVCAR3 ovarian cancer cells were seeded in the assembloid core with NK cells in the collagen matrix. Scale bar, 500 μm.
